# Inhibitory Control, Shifting, and Working Memory Updating Domains form Cognitive Phenotypes in Non-human Primates

**DOI:** 10.64898/2026.02.24.707708

**Authors:** Xuan Wen, Leo Malchin, Adam Neumann, Thilo Womelsdorf

**Author notes:** **Corresponding Authors**: Dr. Thilo Womelsdorf, Vanderbilt University, Psychology Department, 301 Wilson Hall, 111 21st Avenue South, 37240-1103 Nashville TN.

## Abstract

Executive functions comprise at least four major subdomains: Inhibitory Control, Updating, Shifting, and Working Memory. Cognitive abilities in these subdomains are partially separable and partially unified in a common cognitive control factor in humans, but how these functions are organized in the nonhuman primate (NHP) is largely unknown. Here, we used a multi-task assessment approach and found that NHPs show within single sessions reliable cognitive markers of Inhibitory Control in an antisaccade task, (*ii*) Updating abilities in a multidimensional continuous updating task, (*iii*) Working Memory in a delayed matching task, and (*iv*) Shifting abilities in a feature-based rule learning task. First, we found that subjects’ performance fell into three separable cognitive phenotypes with unique strengths and weaknesses across cognitive subdomains. Second, beyond individual tasks cognitive metrics gave rise to four latent cognitive factors separating abilities for shifting/learning and working memory updating, and distinguishing abilities in inhibitory control of exogeneous versus endogenous interference. These results document four separable latent cognitive factors underlying executive control performance in this NHP sample, with inter-subject differences of these factors forming cognitive phenotypes.

## Introduction

Executive functions (EFs) describe higher-order, domain-general cognitive abilities that have a pervasive influence on coping with environmental demands(*1*). But how EF subfunctions are organized in nonhumane primates is largely unknown and indirectly inferred from neuroscience studies of the prefrontal cortex(*2*). In humans, cognitive studies have distinguishes four broader cognitive EF constructs: Shifting, Inhibitory control, Updating and Working memory(*3*). These constructs describe domain-general abilities that have trait-like stability within individuals but vary between individuals, predicting how well subjects use the EFs across modalities and tasks(*3, 4*). Beyond the separability – or diversity – of these four domains, large-scale cognitive studies have identified a common EF factor that has been labeled attentional control, inhibitory control, or cognitive control(*5–7*). This common EF factor describes latent, shared variance of performance across tasks that have high demand on shifting, inhibitory control, updating and working memory. There is ample support for a shared cognitive resource in humans supporting common EF functions including brain network nodes activated in a domain-general way across tasks(*8, 9*), common alterations of large-scale brain networks associated with EF dysfunctions in psychopathologies(*10*), a pool of genes associated with a unified EF factor(*11, 12*), and evolutionary pressures that may not only have shaped modular adaptations for coping with specific demands of an ecological niche but that also will have favored a more unified, general problem solving EF ability(*13–16*).

The rich insights into a cognitive architecture of separable and unified EF functions in humans contrast to the few insights about the relationship of higher cognitive functions in nonhuman primate species. While there is early evidence for a latent common cognitive factor underlying reasoning and tool use, as well as for a shared spatial cognitive ability in great apes(*17*), studies comparing multiple cognitive abilities among nonhuman primate species have found rather independent, uncorrelated abilities across tasks with varying attentional control demands(*18, 19*), or they reported selective correlations of cognitive subdomains such as working memory and flexible sequential reasoning abilities(*20*).

One reason for the scarcity of insights into the cognitive organization of executive functions in NHPs is a lack of assessment batteries allowing to train and assess nonhuman primates on multiple higher cognitive domains, which is surmounted only recently by touch-screen based task batteries(*19, 21–25*). Another reason for lacking insights into the cognitive organization of EFs in NHP’s is the historical emphasis of neuroscience studies to use variations of single tasks to identify functionally specific contributions of neuroanatomical sub-regions of the prefrontal cortex and connected areas(*2, 26–30*). Only recent studies have started to evaluate how separable cognitive functions recruit shared neuronal resources in NHP brain networks(*31, 32*), but also how executive functions may be realized by less integrated neuroanatomical connectivity in NHP compared to humans(*33*).

To close our gap in understanding the organization of EFs in NHP we set out to characterize the cognitive abilities in tasks that have high loading on the constructs shifting, inhibitory control, updating and working memory. We assessed these constructs in six rhesus monkeys performing four tasks within ∼2h repeatedly within 26-58 sessions. This multi-task assessment revealed cognitive phenotypes showing either superior general performance across all tasks, or specific weaknesses in inhibitory control or working memory and set shifting. Beyond these subject-specific cognitive profiles we found latent cognitive factors using exploratory factor analysis that distinguished working memory updating, shift/learning, as well as two separable abilities of inhibitory control of interference of mental (endogenous) versus bottom-up (exogeneous) sources. These findings establish a reference of a possible 4-factor architecture of executive functions in NHP’s.

## Results

We assessed the constructs inhibitory control, shifting, updating and working memory with four different tasks performed in a fixed temporal order in each experimental session: A Delayed Match-to-Sample (*DMTS*) task for working memory (WM), a Continuous Updating (*CU*) task for updating and WM, an accuracy-based Antisaccade (*AS*) task for inhibitory control, and a Flexible Learning (*FL*) task measuring feature-rule learning and set shifting (**Fig. 1**). We adopted these tasks because they prominently recruit the cognitive constructs. Six subjects performed all four tasks in 264 sessions (subjects B: 44; F: 49; I: 44; R: 43; S: 26; W: 58), each lasting on average ∼113.88 (SE: 1.27) min. Subjects self-initiated the tasks on touchscreen Kiosk stations in their home cages running the multi-task software platform M-USE(*24, 34*). Example performances of each task are shown in **Movies’s S1-4**.

**Fig. 1.**
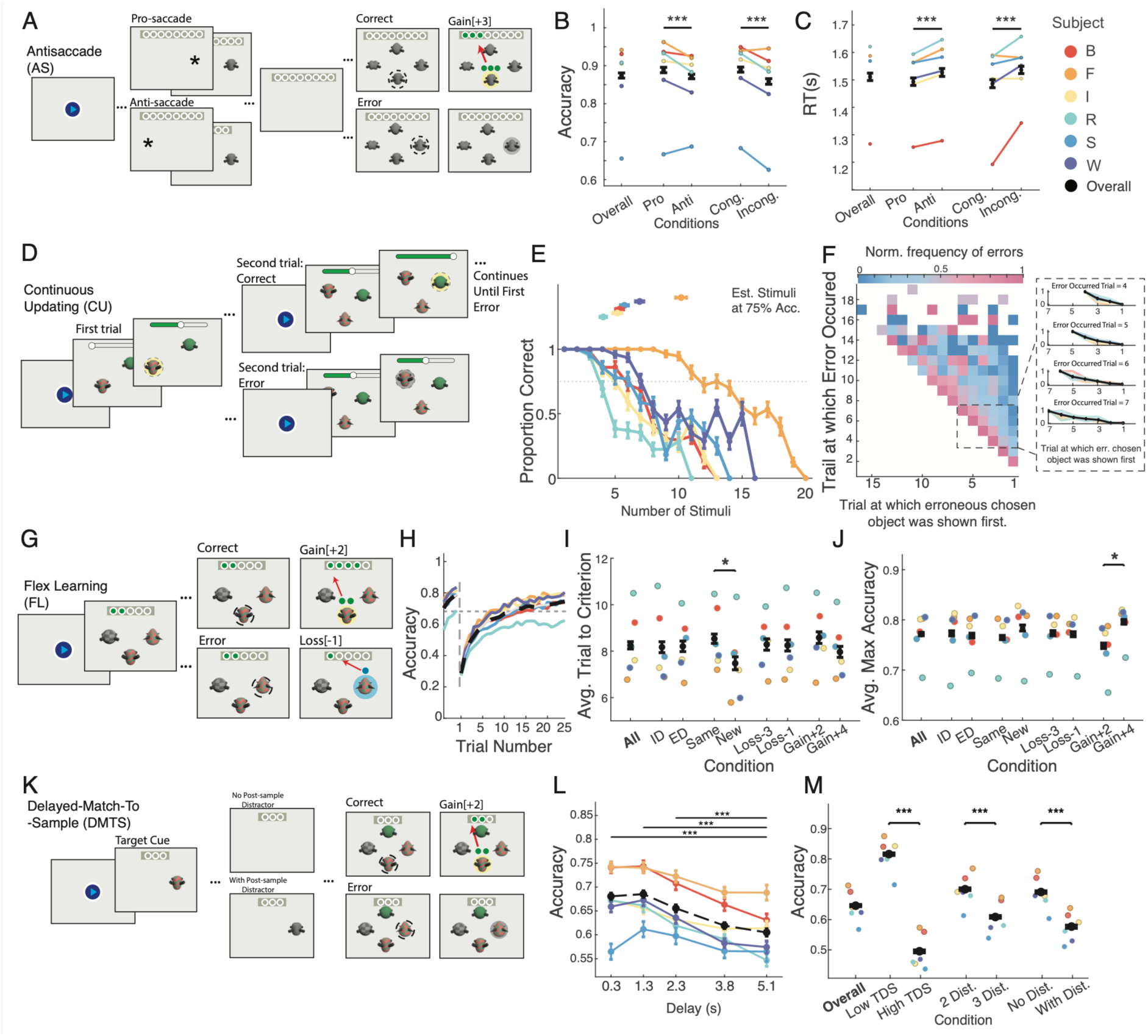
Task paradigms and behavioral results. (**A**) Trials of the Antisaccade (AS) task. Subjects fixate on a central point, a cue appears peripherally, and a target object is briefly presented either at the cue’s location (Pro-saccade) or the opposite location (Anti-saccade). After a brief delay four objects appear as response options and subjects have to select the target object to receive +3 tokens. (**B**) Accuracy across AS task conditions. *** indicate p<0.001 (pairwise t-test). (**C**) Same as *B* for reaction times. (**D**) Example trials of the Continuous Updating (CU) task. The first trial presents two objects; each subsequent trial adds one object; Subjects have to choose an object they haven’t previously chosen to receive reward. (**E**) Prop. correct with increasing number of stimuli in the display (*x-axis*). Colored dots show mean (±SE) number of stimuli at which subjects were correct in 75% of blocks. (**F**) Norm. frequency that the erroneous choice was for an object introduced n trials before the erroneous trial. The inset visualizes that errors were most likely for objects chosen earliest in a trial and least likely for more recently chosen objects. (**G**) The Flexible Learning (FL) task showed 3 objects. Subjects chose one object to receive tokens, learning by trial-and-error which object feature yields tokens. (**H**) Accuracy drops when the rewarded object feature changes and recovers with learning (colored lines: subjects). (**I**) Avg. FL trials-to-criterion for 8 block conditions. A * indicates p<0.05 significance (Bonferroni corrected). (**J**) Avg. FL plateau accuracy across conditions. (**K**) Delayed Match-To-Sample (DMTS) task showed a sample object, delay, and a response display requiring subjects to choose the object matching the sample. (**L**) Accuracy over delays. *** indicates p<0.001 (Bonferroni corrected). (**M**) DMTS accuracy at low/high Target-Distractor similarity (TDS); when response displays had 2 or 3 distractor objects, and with/without a distractor flashing during the delay. Colors represent individual subjects, black represents overall average (± SE).

The *Antisaccade* (*AS*) task assessed inhibitory control abilities by presenting a brief target stimulus either at the same (pro-) or opposite (anti-) location relative to a brief attention-grabbing peripheral cue (**Fig. 1A**). The target was followed by a response display with four objects. Subjects had to recognize the target in the response display, which makes this task an accuracy-based AS task that we adopted from and named according to prior studies(*6*). In contrast, to the original AS task(*35*), the accuracy-based AS task operationalized control of covert attentional shifting to the target object as opposed to inhibition of an overt eye movement. Subjects showed lower accuracy and slower reaction times on anti-than prosaccade trials (prop. correct difference pro- vs antisaccade: 0.017±0.002, p<0.001; and for reaction times: 34 ms ±6, p<0.001) (**Fig. 1B**,**C**,**S1A-C**). Accuracy and reaction times were also lower in trials in which the target object in the response display was presented at an opposite (incongruent) location when compared to the same (congruent) same side as the preceding target location, which signifies the so-called spatial congruency Simon effect (prop. correct difference congruent vs incongruent: 0.031±0.004, p<0.001; reaction times: 50 ms ±7 ms, p<0.001) (**Fig. 1B,C**). Both, the AS effect and the Simon effect were consistently evident in individual sessions (**Fig. S1A-F**). The *Continuous Updating* (*CU*) task assessed WM updating abilities by asking subjects to choose one among many objects in a display that they have not chosen before. Consecutive trials show all previously chosen objects at random locations plus a new object. To avoid choosing an object twice it needed to be updated in WM once chosen (**Fig. 1D**). Subjects reached on average 6.30 objects (SE 0.18; range: 4.47−10.18) before they committed an error with a p=0.75 probability (quantified with the metric: *CU_Accuracy_TrialAt75Acc*), which indexes the WM updating ability and is a measure of WM capacity (**Fig. 1E**). When subjects the trial and least likely objects chosen in the most recent, previous trial. This effect was quantified with the metric *CU_nBack_Slope* whose positive slope (on average 0.24 ±0.02, p < 0.001) signifies temporally more remotely chosen objects were more likely erroneously chosen than more recently chosen objects (**Fig. 1F, S1G**). The reaction times in the CU tasks systematically slowed down with increasing number of objects in the display, which shows that subjects searched the increasing object set before committing to a choice (avg. reaction time set size slope: 0.082±0.006, p<0.001) (**Fig. S1H-J**).

The next *Flexible Learning* (*FL*) task assessed the ability to learn and shift attention sets. It measured how many trials subjects needed to learn through trial-and-error which object feature was associated with reward. Each trial presented three objects that differed in six feature values of two feature dimensions (e.g. three shapes and three colors) (**Fig. 1G**). One feature value (e.g. an oblong shape) was designated the rewarded target for 25-37 trials and was either from the same or a different feature dimension as the target in the preceding block of trials, which signifies intra-versus extra-dimensional shifts (conditions: ID, ED) of the target feature. A block of trials either used the same features as in the previous block or it used novel features that were not shown before (conditions: Same, New), it rewarded correct trials with few or many tokens (conditions: Gains +2, +4) and penalized error trials with the loss of one or three tokens (conditions: Loss -1, -3) (**Fig. 1G**). Subjects learned, i.e. reached the performance criterion, within 8.25 trials in a block (range: 6.78–10.49 trials) and reached on average 77.18% plateau accuracy (range: 68.48%–80.54%) (**Fig. 1H**). Learning was faster when the target feature was new as opposed to a feature that was previously a distractor (trials-to-criterion: New: 7.48, Same: 8.54; p<0.05) (**Fig. 1I**). In blocks with a higher incentive accuracy was higher (G4: 0.796±0.005 vs. G2: 0.748±0.005, p<0.05) (**Fig. 1J**), and perseverative errors were reduced (G4: 0.059±0.001 vs. G2: 0.073±0.001, p<0.05) (**Fig. S2A**).

The fourth Delayed Match-to-Sample (DMTS) task assessed WM by tracking how accurately a sample target was recognized within a response display after 0.3-5.1 s delays (**Fig. 1G**). Performance decreased with increasing delay (**Fig. 1L**), evident in a negative regression slope of accuracy over delay (metric *DMTS_Accuracy_Slope*: r=−0.020±0.003, p<0.01) (**Suppl. Fig. S2D**). DMTS performance was lower when the target-distractor similarity (TDS) in the response display was higher (High TDS: 0.487±0.005 vs. Low TDS: 0.807±0.004; p<0.001), when the response display included three rather than two distractors (3 Distr: 0.600±0.004 vs. 2 Distr: 0.692±0.004; p<0.001), and when a distractor object was briefly flashed during the delay period (With distractor: 0.568±0.005 vs. No distractor: 0.682±0.004; p<0.001) (see: **Fig. 1M**). This result pattern is consistent with human studies (e.g.(*36, 37*)). A similar pattern was observed for reaction time variations across task conditions (**Fig. S2D-G**).

The four tasks allowed operationalizing 39 cognitive metrics of whom 29 showed high (≥0.5) re-test reliability across sessions and were used for further analysis (**Fig. 2A-C**) (for metrics definitions, *see* Methods). Metrics based on choice accuracy had re-test reliability scores of ≥0.96 while metrics based on reaction times had reliability scores of 0.87-0.98 (**Fig. 2C**). The tasks provided 16 metrics with >0.5 reliability scores indexing inhibitory control of different types of interference (**Fig. 2B**). Six metrics indexed motivational effects of variable gains and losses on FL task performance, but these metrics showed low reliabilities (≤0.21) across sessions (**Fig. 2B**,**C**). The exception was a systematic reduction of perseverative choices in the FL task in blocks when errors threatened the loss of 3 compared to 1 reward token (reliability score of metric *FL_CEn_Perseveration_Loss_Index(i)* = 0.83), which signifies that anticipating larger losses increased motivational effort control(*38, 39*).

**Fig. 2.**
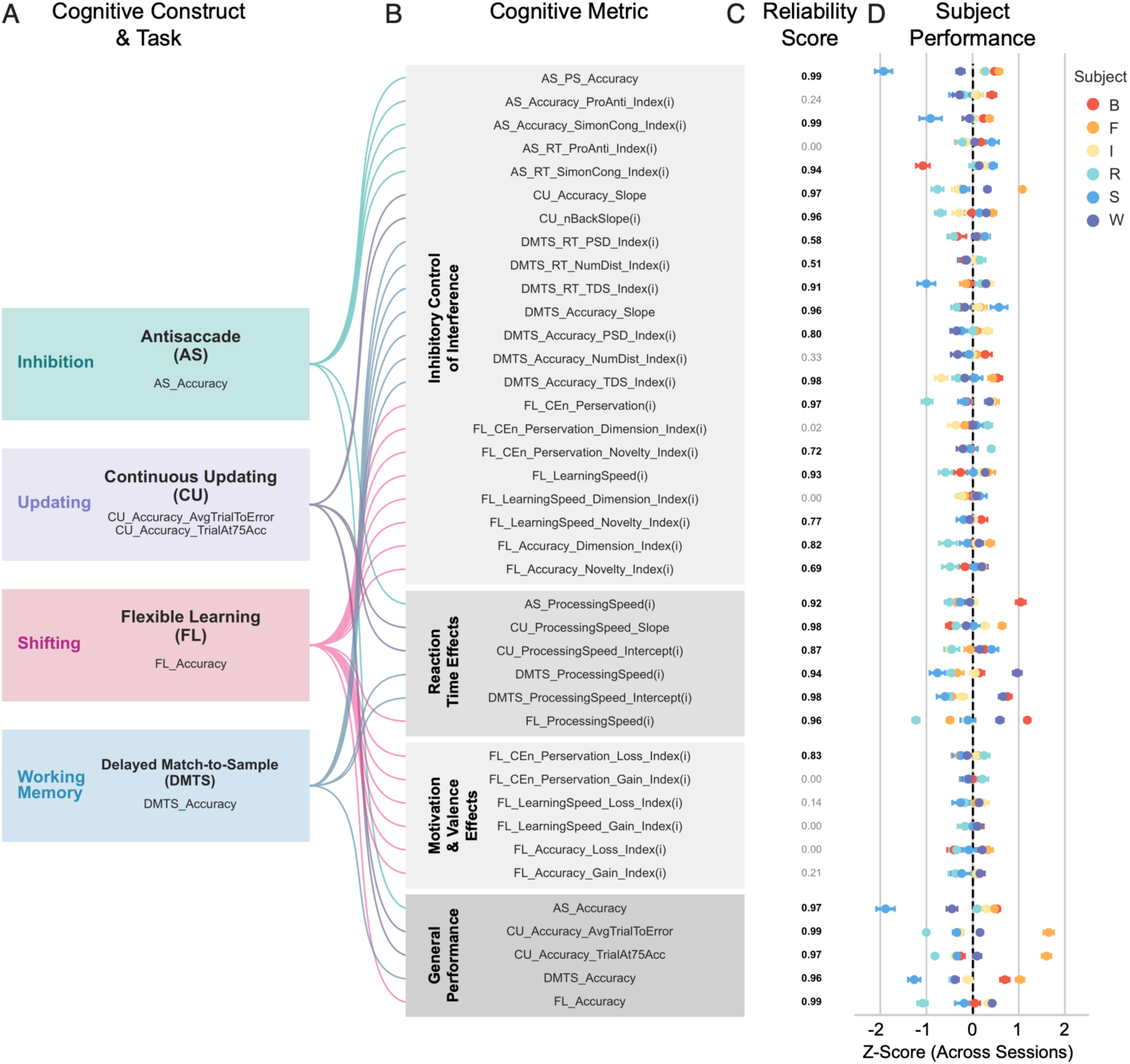
Cognitive profiles across task metrics and their re-test reliability. (**A**) Four tasks in colored boxes and the main cognitive domain they recruit. (**B**) Metrics derived from all tasks, grouped into Inhibitory Control, Processing Speed, Valence, and General Performance. Metrics appended with ‘(i)’ are sign-inverted such that a higher value consistently indicates better performance (i.e. less interference from distractors). Metrics ending in ‘*_Index*’ represent the norm. difference between two conditions. (**C**) Spearman-Brown split-half Reliability Scores. Scores > 0.5 are in bold font to indicates they passed the reliability criterion. (**D**) Performance Z-scores (± SE across sessions) for each subject (in different colors). The dashed vertical line indicates the group mean (Z=0).

### Subjects are grouped into three cognitive phenotypes

To compare subjects’ performance, we z-standardized the performance of each cognitive metric. The resulting cognitive profile shows that individual subjects could under- and over-perform relative to the group mean by up to -1.89 (subject S) to +1.66 (subject F) z-normalized STD values (**Fig. 2D**, **Suppl. Fig. S3A**). Within cognitive domains, subjects varied from the group mean in a similar range for inhibitory control (minimum and maximum difference to group mean: z = -2.0654 (S) to 0.8166 (B)), set shifting (z = -2.0433 (R) to 0.8277 (W)), continuous updating (z = -1.0372 (R) to 2.0986 (F)) and working memory performance (z = -1.5754 (S) to 1.4518 (F)) (**Suppl. Fig. S3B**). To discern underlying patterns of this performance variability we used dimensionality reduction across 29 metrics of the four domains and identified clusters based on the two main dimensions. This separated three clusters containing scores mostly of subjects R and I in cluster 1, scores from subjects B, S, and W in cluster 2, and scores of subject F in cluster 3 (**Fig. 3A**, **B**). The grouping of subjects suggests that clusters were stable across sessions forming cognitive phenotypes. Confirming this impression, we found subjects I and R were clustered into phenotype 1 in on average 93.9% of sessions; B, S, and W were clustered into phenotype 2 on average in 87.7% of sessions, and subjects F constituted phenotype 3 in 89.6% of experimental sessions (**Fig. 3C**). To interpret the cognitive phenotypes we extracted the performance scores of the four tasks, which showed that phenotype 1 (subjects I and R) had z-scores indicating relatively poor performance of the FL (*Shifting*, z = -1.32 ± 0.45), DMTS (*WM*, z = -0.83 ± 0.11) and CU (*Updating*, z = -0.97 ± 0.07) task, while phenotype 2 (subjects B, S, and W) showed relatively poor scores of the AS task (*Inhibitory Control*, z = -1.10 ± 0.30) (**Fig. 3D**). In contrast, phenotype 3 (subject F) exceeded performance with z values >1 in all four tasks.

**Fig. 3.**
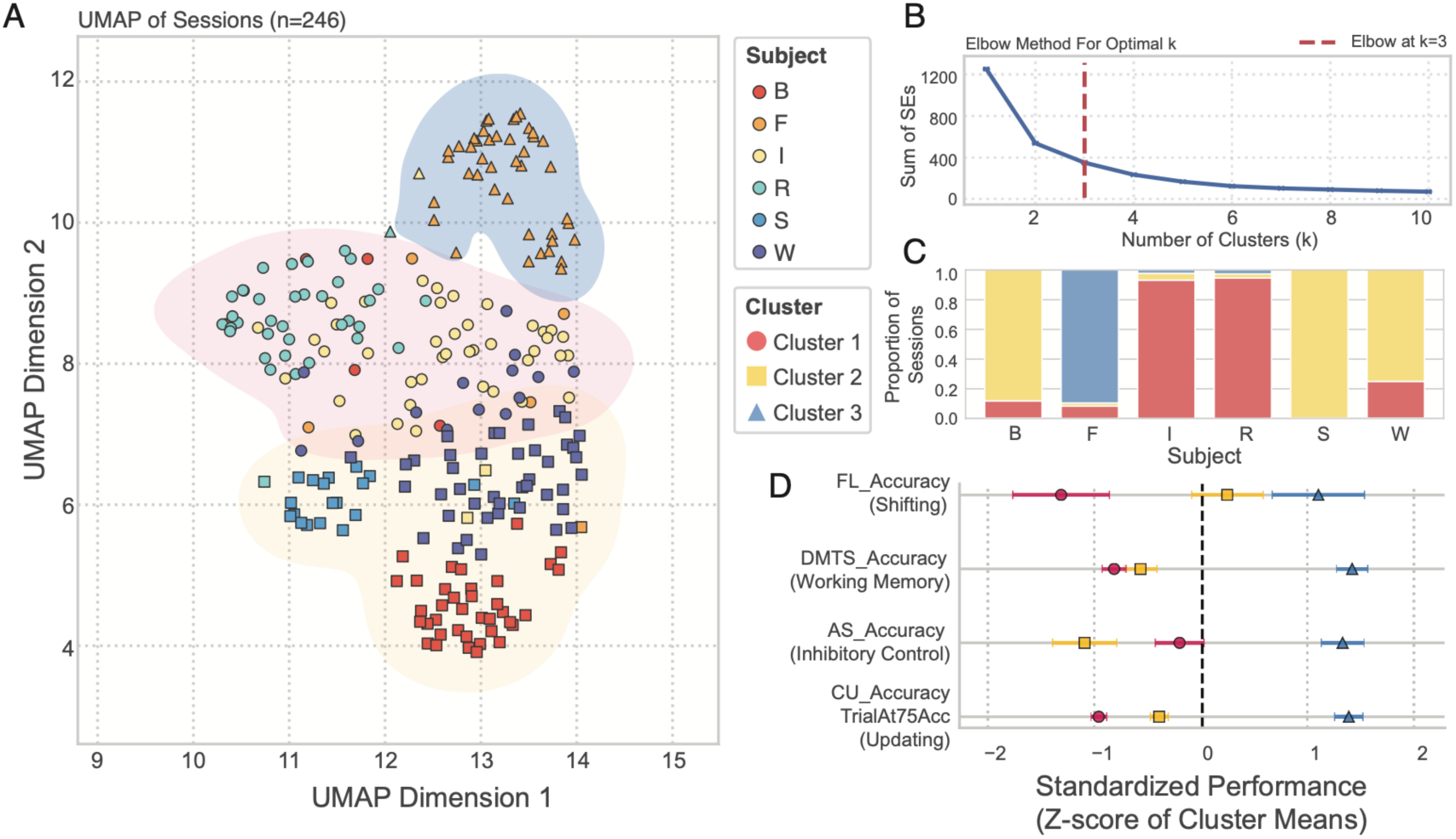
Unsupervised Discovery of Cognitive Phenotypes. (**A**) Uniform Manifold Approximation and Projection (UMAP) embedding of all sessions (n=246) based on z-scored performance metrics. Each point represents a single session. Point color indicates the subject; symbols and shaded contours represent the k-means cluster ID. K-Means was applied to the 2D UMAP coordinates. (**B**) Sum of squared errors (SEs) for a range of cluster numbers (k). The dashed red line at k=3 indicates the selected optimal number of clusters (based on the Elbow method). (**C**) Stacked bar chart for each subject (*x-axis*) showing the proportion of sessions the subject was assigned to one of the three clusters. The high consistency of cluster assignment suggests clusters represent cognitive phenotypes. (**D**) Standardized performance profiles for each cluster. Points represent the mean Z-score (± SE) for each cluster on the four primary task performance metrics. The dashed vertical line indicates the population grand mean (Z=0).

### Positive metrics-level correlations across cognitive domains

While subjects of the same cognitive phenotype show similar performance across metrics, there might also be shared variability of metrics beyond individual subjects. We tested this by calculating partial correlations across the 22 reliable behavioral metrics (**Fig. 4A**). These metrics-level correlations were on average positive but weak (r=0.064, p<0.05) (**Fig. 4B**). Correlations remained significant when only metrics from different tasks were considered (between-task correlations: r = 0.0238 ± 0.0089, p = 0.00903, n = 80), were stronger for metrics from the same task (avg. within-task metric correlations: r = 0.1620 ± 0.0542, p = 0.00728, n = 25) and when considering only reaction time metrics (**Suppl. Fig. S4**).

**Fig. 4.**
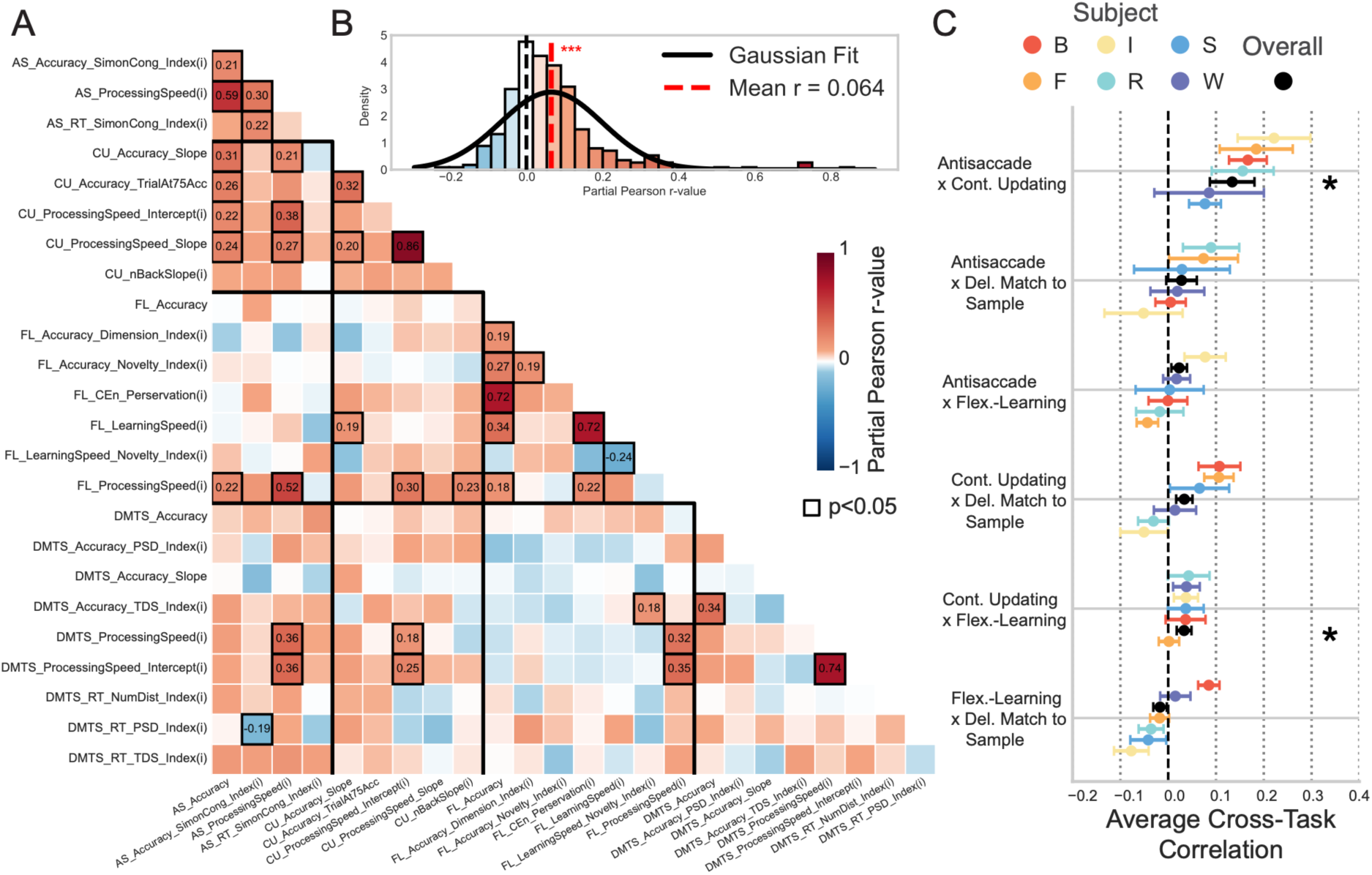
Correlations between metrics and tasks. (**A**) Partial Pearson r-values for all behavioral metrics that passed the reliability threshold. Metrics are ordered and grouped by task (AS, CU, FL, DMTS), with heavy black lines separating task blocks. Significant correlations (p < 0.05, FDR corrected) have black border and their r-value as text inside the cell. (**B**) Distributions of correlations from panel *A*, overlaid with a fitted Gaussian. Red vertical line marks the mean correlation (r = 0.064) which was different from 0 (one-sample t-test, p < 0.001), indicating a low positive correlation of metrics. (**C**) Cross-task correlation (mean r-value ± SE) for each subject (colors) and overall (black). Values represent the mean partial correlation computed between all reliable accuracy-based metrics from the first task or a pair and all reliable accuracy-based metrics from the second task. Asterisks (*) indicate pairs with an overall mean correlation different from zero (p < 0.05).

Focusing on between-task correlations we found that metrics of the updating (CU) task correlated with inhibitory control metrics of the AS task (r = 0.1339 ± 0.0466, p = 0.0365), with all six subjects showing positive correlations (**Fig. 4C**). In addition, metrics indexing updating performance (CU task metrics) correlated weakly with metrics indexing shifting performance (task: Flex-Learning) (r = 0.0333 ± 0.0155, p = 0.0463), with five of six subjects showing low, but positive correlations (**Fig. 4C**).

### Latent cognitive factors distinguish set shifting, updating and inhibitory control abilities

Correlations of metrics-level and task-level performance scores are positive, but weak not only in in our study (**Fig. 4**), but also in human cognitive studies with r’s in the range of ∼0 - 0.17(*7*). Stronger relationships have emerged in latent factor analysis. Our dataset satisfied the criteria for an exploratory factorial (EF) analysis(*40*), which we applied to the fourteen accuracy-based metrics, similar to previous studies(*7, 41, 42*). The EF analysis identified four separable latent cognitive factors (Kaiser’s criterion Eigenvalues > 1) (**Suppl. Fig. S5A**). The four-factorial structure was influenced by both, between-subject differences across sessions and within-subject variance, but was evident considering each source of variability separately (**Suppl. Results** and **Suppl. Fig. S6**). Each of the four factors contained 3-4 metrics with higher positive factor loadings. The first factor was composed of high loadings of metrics from the FL task that signified (*i*) faster set shifting and learning speed (*FL_LearninSpeed(i)*), (*ii*) less perseverative choices (*FL_CEn_Perseveration*), and (*iii*) higher overall set shifting accuracy (*FL_Accuracy*) (**Fig. 5A**). These metrics suggest labeling factor 1 *Shifting/Learning*. The second factor showed high loadings of metrics from three tasks, grouping together performance that signified less interference from target-distractor similarity in the DMTS task (*DMTS_Accuracy_TDS_Index(i)*), higher antisaccade accuracy (*AS_Accuracy*), less interference from spatial incongruency (i.e. less Simon effect, *AS_Accuracy_SimonCong_Index(i)*), overall higher DMTS accuracy (*DMTS_Accuracy*), and a moderate, positive loading of the updating performance in the CU task (*CU_Accuracy_TrialAt75Acc*) (**Fig. 5A**). High positive factor loadings of these metrics mostly signify better ability to control interference from external sources that are based on relations of the stimulus features and spatial locations, suggesting as a tentative label ‘exogeneous Inhibitory Control’.

**Fig. 5.**
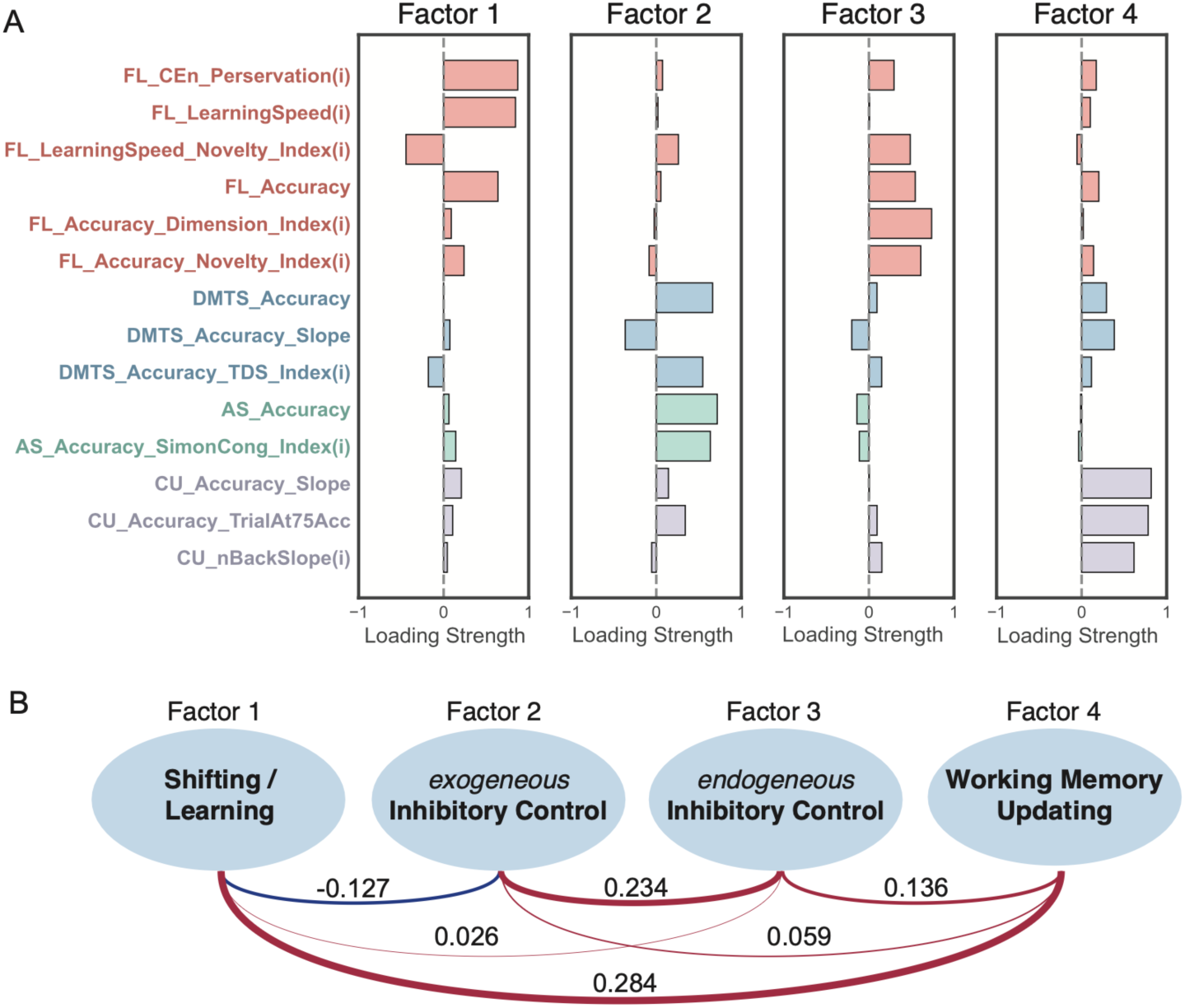
Latent factor structure underlying metrics scores. (**A**) Loadings (*x-axis*) of each factor of the 4-factor solution across behavioral metrics (*y-axis rows*). The 4-factor solution was determined using the Kaiser criterion (Eigenvalue = 1); factor loadings were derived using principal axis factoring with a Varimax rotation. Metrics are color-coded by their task domain. (**B**) A diagram of the inferred latent factor structure. The four factors were interpreted and named based on the high-loading metrics shown in *A*. The lines connecting the factors illustrate their pairwise correlations from the exploratory factor analysis. The numerical value, line color (Red=positive, Blue=negative), and line width represent the strength and direction of the correlation.

The third factor showed high loadings of FL-Learning metrics that did not load on factor 1 and which quantified differences in learning performance between learning conditions: It entailed metrics that contrasted how strongly accuracy and learning was affected by whether the target feature had a history as distractor in the previous learning block (New vs Same condition, metric *FL_LearningSpeed_Novelty_Index(i)*), and whether learning involved extra-versus intra-dimensional target shifts (*FL_Accuracy_Dimension_Index(i)*) (**Fig. 5A**). These metrics index how flexible subjects adjust internal top-down target representations which we summarize as reflecting inhibitory control of endogenous representations. This interpretation contrasts with factor 2 for which inhibitory control requirements were based on external stimulus features (target-distractor similarity) and stimulus locations (antisaccade and spatial congruency Simon effect). To address the difference to factor 2 we label factor 3 ‘endogenous Inhibitory Control’. Factors 1-3 differed from factor 4, which had high loadings of metrics from the CU and DMTS tasks signifying the updating of working memory (*CU_Accuracy_Slope*, *CU_Accuracy_TrialAt75Acc*), and more stable accuracy across delay durations (*DMTS_Accuracy_Slope*, for which a higher loading corresponds to a shallower slope of accuracy across delays, i.e. less performance decline with increasing delay) (**Fig. 5A**).

To test the robustness of the factorial analysis results, we used the same metrics and applied a principal component analysis with subsequent hierarchical clustering. This approach separated metrics into four clusters that grouped together metrics similar to the grouping of the four factors of the EF analysis (**Suppl. Fig. S5B,C**). Clusters signifying working memory updating, shifting/learning and endogenous inhibitory control contained the same metrics as the corresponding EF factors, while the fourth PCA-based cluster resembled the endogenous inhibitory control EF factor but included two metrics that had no apparent loading on EF factor 3, indexing the decline of accuracy with increasing delays in the DMTS task (*DMTS_Accuracy_Slope*) and the bias of CU task to commit errors more likely of objects the longer they were retained in WM (*CU_nBackSlope(i)*) (**Suppl. Fig. S5C**).

### No evidence for a common factor, but selective correlations among the four cognitive factors

We next tested whether the 4-factor solution of the exploratory factor analysis is a good description of the data when compared to model-based confirmatory factor analysis (CFA), bifactor analysis that assumed one common general factor, and higher-order general-factor analysis (*see* **Method**s). First, we found that a CFA supported the 4-factorial structure of the exploratory factor analysis model, showing a superior fit to the data than other factor structures, although the overall fit quality was modest (**Suppl. Fig S7A-D**). Second, we found no evidence for a shared common factor. Hierarchical CFA, different bi-factorial structures and higher-order analysis that assumed a common general factor showed low reliability and consistently scored below 4-factor solutions (**Suppl. Fig. S7C,E,F**; **Suppl. Table S1-6**). We also tested whether the inclusion of reaction time metrics into latent factor analysis would provide evidence for a general factor but found no evidence that variations of processing speed / reaction contribute as a general factor (**Suppl. Results**). Taken together these null findings quantify that in our sample of six subjects and across metrics from four tasks subjects’ abilities are not accounted for by a shared factor.

While there was no evidence for a single shared factor, we tested for more specific correlations among the four latent factors that were separable using EFA and PCA-based clustering. We found that exogenous and endogenous Inhibitory Control factors (EF factors 2 and 3) correlated at the latent factor level (r=0.234; p<0.05) (**Fig. 5B**). A similar correlation value (r=0.284; p<0.05) was evident for Shifting/Learning (EF factor 1) and Working Memory Updating (EF factor 4). Endogenous Inhibitory Control (factor 3) and Working Memory Updating (factor 4) correlated positively but only weakly (r=0.136; n.s.). These three positive latent correlations contrasted to three near-zero correlations. Factor 1 (Shifting/Learning) was uncorrelated with both Inhibitory Control factors 2 and 3 (r = -0.127 and r = 0.026; both n.s.), and endogenous Inhibitory Control (factor 2) was unrelated to Working memory Updating (factor 4) (r = 0.059; n.s.) (**Fig. 5B**).

We interpreted EF factors 2 and 3 as reflecting different types of inhibitory control abilities. The latent factor correlation (r=0.234) of these factors may point to interdependencies of performance scores at the level of the metrics defining these factors. To test this possibility, we evaluated metric-level correlations of all those performance scores that operationalized inhibitory control of interference. This included metrics contrasting performance (***1***) with increasing number of distracting objects in the display (*DMTS_Accuracy_PSD_index(i)*, *CU_Accuracy_Slope*), (***2***) with increasing perceptual interference due to the number of shared features of distractor and target objects (*DMTS_Accuracy_TDS_index(i))*, (***3***) with interference from distractor objects because they were previously targets as opposed to new, unfamiliar distractor objects (*FL_Accuracy_Novelty_Index(i)*), (***4***) with interference from distracting features of the same versus a different feature dimension than previous targets (*FL_Accuracy_Dimension_Index(i)*), and (***5***) with interference from distractor objects appearing in the response display of the antisaccade task on the opposite side as the target stimulus (*AS_Accuracy_SimonCong_Index(i)*). We found that correlations among these inhibitory control metrics were low and not different from zero (avg. r = 0.022 ± 0.020, p = 0.2913) (**Suppl. Fig. S8**). Thus, different types of inhibitory control showed no overall task-level correlation, while analysis of shared latent correlations separated two distinguishable types of inhibitory control abilities and revealed a moderate latent correlation among these factors signifying exogenous and endogenous inhibitory control.

## Discussion

We have shown that NHP’s provide reliable performance scores across multiple cognitive metrics of a 4-task-per-session assessment assay (**Fig 1**,**2**). The scores grouped the six individual subjects of our study into three cognitive phenotypes that signified (*i*) selective weaknesses in inhibitory control, or (*ii*) in shifting/learning and working memory updating abilities, or (*iii*) that showed superior performance across all task domains (**Fig 3**). Beyond individual cognitive profiles, the metric scores gave rise to four latent cognitive factors that suggest separable latent variance underlying shifting/learning, working memory updating and inhibitory control of interference from exogeneous and endogenous sources (**Fig. 5**). These results support a 4-factor model of executive functions in our sample of six subjects and provide a strong quantitative prediction that a similar higher-order cognitive structure will be evident in larger samples of nonhuman primates. The pattern of results emphasizes the relative independence of four separable cognitive factors and how individual variations of them form cognitive phenotypes.

Illustrating the separability of executive sub-domains is a major finding because the data could have supported alternatives such as a stronger positive manifold among performance scores, or evidence for a common EF factor(*43*). In contrast, we found overall low correlations among performance scores (**Fig. 4**, **S8**) and robust statistical uniqueness of latent cognitive factors with rather weak shared variability among specific executive control functions (**Fig. 5**, **S7**). Taken together, these results provide a rich multi-task characterization of EFs in NHP and extend the literature on the unity and diversity of executive cognitive functions that so far is based on humans and on few studies with great apes.

### Performance profiles suggest cognitive phenotypes

Performance scores across the four tasks grouped subjects into three clusters. We interpreted these clusters as cognitive phenotypes to highlight that there is a consistent structure of individual differences in executive functions in our sample of subjects, similar to prior cognitive phenotyping studies(*15, 44*). We consider the evidence for three cognitive phenotypes as an important proof of principle that cognitive profiles of individual subjects do not randomly vary across individuals when evaluated over multiple sessions but rather show meaningful subject-specific patterns that deviate from what is expected from a broad normal distribution. Proving the robustness of such cognitive phenotypes will require larger sample sizes and reliable multi-task assessments. So far, extended measurements of multiple higher cognitive abilities have been rare in nonhuman primates(*21–23*), but emerging evidence in great apes that suggests that higher cognitive abilities can show trait-like temporal stability(*17, 45, 46*). Among the three cognitive phenotypes we found, one subject stood out to show superior performance in all four tasks compared to all other subjects (cluster 3, monkey F, **Fig. 3**). This subject may benefit from some common neuronal resource shared across executive function domains. This result warrants future studies to search for the neurophysiological and genetic origins of a domain-general, common executive control ability(*12, 47*). Another phenotype (cluster 1, subjects I+R) scored one STD below the average in the working memory updating and set shifting task. Both tasks require the maintenance and updating of an internal ‘top-down’ goal representation over multiple trials, which was an object-based representations in the CU task, and a feature-based rule representation in the FL task. Common demands for mental maintenance and updating are consistent with the statistically significant correlation of the latent cognitive factors Shifting/Learning and Working Memory updating (**Fig. 5B**). The last phenotype (cluster 3, subjects B+S) showed below one STD performance only in the AS task, which indexes difficulty inhibiting reflexive orienting towards a salient cue. This finding suggests that inhibitory control deficits may relate to previously described phenotypes of impulsivity which characterize humans with substance use disorder and obsessive-compulsive symptomatology(*48, 49*). Selectively compromised inhibitory control performance of the AS task also suggests individual difference of overall attentional control abilities because human studies found that the accuracy-based AS task, as deployed here, has the highest loading among diverse tasks on a common attentional control factor compared with other cognitive control tasks(*42*). Consistent with such a more general attentional control effect, the subjects with larger z-score deviations in AS task performance also had relatively poorer DMTS and CU working memory performance scores (**Fig. 3**). It will be important to understand in future studies whether subjects of the same cognitive phenotype with lower-than-average performance in subdomains of executive functions benefit more prominently than subjects of other phenotypes from neuropharmacological interventions targeting those domains(*26*).

### Working memory updating is distinguishable from inhibitory control

Our results distinguished a latent factor of Working Memory Updating measured with the CU task and factors indexing Inhibitory Control of interference and flexible Shifting/Learning (**Fig. 5**). The separability of a unique Working Memory Updating factor is reminiscent of individual difference studies in humans that have distinguished a working memory factor from an inhibition/shifting related cognitive factor(*7*). In humans, the separation of these latent factors emerges during development around ∼6-10 years of age(*50–52*). The core cognitive demands of the CU task that underlie the Working Memory Updating factor involved maintaining an increasing number of objects in short-term memory and updating it with novel items. These demands are similar to those of forward/backwards span tasks used in humans. The working memory span tasks are sometimes labeled short-term memory tasks in human studies because they lack additional attention-demanding operations such as solving math equations during the delay while keeping track of a growing number of items in short-term memory (e.g.(*53*)). The distinction of a short-term memory and a working memory with an attentional control component has an important history that either considered the attention component a ‘*central executive*’ separated from short-term maintenance(*54, 55*), or that suggest that working memory is integrated with attention as in the ‘*focus-of-attention*’ WM model(*56*). These considerations are meaningful because overall fluid intelligence measures in humans using, e.g. the Ravens matrices, correlate stronger with working memory tasks that include attentional control demands than with purer short-term memory tasks(*57*). In our behavioral assay, attentional control demands most closely correspond to inhibitory control of endogenous interference (EF factor 3). Thus, in our sample of NHP’s working memory updating and ‘attentional’ inhibitory control was separable at the latent factor level. One possibility is that attentional control and working memory updating processes are less integrated in NHPs than in humans. Clarifying the precise relationship of working memory and attentional control in NHPs will be an important question for future studies, particularly because of rich evidence for crosstalk between these functions in humans. For example, humans with better working memory show improved antisaccade performance(*58*), make fewer reflexive saccades to the flashing cues in the antisaccade task(*59*), and make fewer errors in Stroop- and Flanker tasks(*60, 61*). These interactions signify that subjects with better WM performance more likely withstand distraction from goal-interfering sensory inputs(*41*). Whether such interactions of working memory and inhibitory ‘attentional’ control processes are evident in cognitive performance of NHP’s will be an important question for future work. Emerging neurophysiological evidence in NHP’s suggest that similar neuronal ensembles in dorsal-lateral prefrontal cortex realize attentional selection and reflect updated working memory representations in NHPs(*31, 32*), while controlling what information is selected and updated may require separate neuronal ensembles in anterior and medial prefrontal brain areas(*62, 63*).

### Interactions of working memory and shifting/learning abilities

Another key finding of our study is the separability of the Working Memory Updating factor from a Shifting/Learning factor (**Fig. 5B**), which resonates with human studies reporting that a Shifting factor is most distinct to factors encompassing working memory or common executive functions, showing sometimes opposing patterns of correlations(*3*). Consistent with these reports we found a trend of negative correlations of Shifting/Learning and the exogenous Inhibitory Control factor (**Fig. 5B**). Such a low anticorrelation may relate to an intrinsic stability-flexibility trade off, which entails that flexibility (supported by stronger Shifting abilities) and stability (supported by stronger Inhibitory Control) are opposing tendencies at the behavioral and neural network level(*5, 64*). Such a trade-off may drive the separability of the cognitive constructs Shifting/Learning and Working Memory Updating at the latent factor level.

At the same time, we found a positive correlation of these factors at the latent factor level (**Fig. 5B**), indicating that working memory updating abilities may also positively contribute to faster learning. This suggestion is supported by studies showing that efficient shifting of attention sets benefits from working memory of recent trials’ choices(*63, 65*). A link of working memory functions with shifting, but not with inhibitory control functions, is also suggested by other behavioral studies in primates. One larger scale behavioral study in chimpanzees found working memory and shifting performance across four tasks were more similar to each other than to inhibitory control performance, measured as the ability to inhibit reaching directly towards a visible-but-inaccessible food in favor of reaching towards an opaque-food loaded cylinder(*66*). The relative independence of set shifting and inhibitory control abilities also has been observed with touchscreen-based assessments in macaques, which found no relationship - indexed as the dissimilarity of subjects’ ranks in different tasks - of reversal learning abilities (number of trials needed to reverse red vs green reward) and the ability of response inhibition in a Go-/No-Go task(*19*). Taken together, our findings and prior literature suggest that shifting and working memory abilities are well separable at the latent factor level, but mutually support each other, while shifting/learning functions are more prominently separable from inhibitory control functions.

### Separable domains for inhibitory control of endogenous vs exogeneous distraction

Our analysis provided two key findings about Inhibitory control. Performance scores of inhibitory control metrics from different tasks were rather uncorrelated (**Suppl. Fig. S8**), while the latent factor level revealed two distinguishable types of inhibitory control (**Fig. 5**). Despite rather moderate loadings, the two factors point to fundamental differences of inhibitory control of interference of endogenous representations (*factor 3*, indexing the effect of DMTS target-distractor similarity) and inhibitory control of interfering exogeneous events such as a salient antisaccade cue and incongruent stimulus-response mappings (*factor 2*, indexing AS task performance and spatial congruency effect in the AS task). We interpret these factor differences as distinguishing the ability to control interference of an endogenous mental representation versus controlling exogenously triggered orienting. Previous latent factor studies in humans have similarly distinguished a factor ‘*interference control from distractors’* as measured with flanker tasks, and a factor ‘*inhibitory control of prepotent responses*’ as measured with AS or Stroop tasks(*67, 68*). These distinct types of interference control are consistent with our findings. In our view, *inhibitory control of distraction* corresponds to controlling an endogenous mental representation, while *inhibiting prepotent responses* is akin to controlling exogeneous spatial orienting.

While there haven’t been previous reports of latent inhibitory control factors in NHP, analysis of task-level analysis points to separable, task-specific inhibitory control abilities. In one elegant set of studies using touchscreen tasks performance was assessed of a Go/No-Go response inhibition task, a distraction task with pictures, and a green/red reversal learning task, and although the relationship amongst these types of inhibitory control measures were not explicitly reported, there were no correlations apparent(*19, 69*). Consistent with this observation, another NHP study reported that inhibitory control abilities in NHPs are task-specific and thus separable(*18, 70*). For example, the ability to inhibit reaching for a remembered previous food location that stopped being baited with food (A-not-B error task) was unrelated to avoiding impulsive reaching to a visible food item (detour cylinder task), or to overcoming a proximity bias in reaching for a food cup in the so-called middle cup task(*18, 70*). Taken together, our finding that inhibitory control abilities are rather unrelated at the task-level but organized into at least two separable latent inhibitory control functions at the latent-factor level provides a framework for future studies to identify the neural basis underlying efficient inhibitory control of interference affecting different types of representations.

### Limitations

Our study documents the separability of latent cognitive factor underlying executive subfunctions in NHP’s based on repeatedly measured, reliable performance metrics using an optimized multi-task assessment battery. Despite the strength of our study the relatively small sample size available for the analysis of task and latent factor correlations constrain the extrapolation of the results to the larger population. This limitation suggests that our study’s strength is to validate a versatile paradigm and provide a strong hypothesis favoring the existence of at least four cognitive factors underlying executive functions. The four-factorial organization of EF’s and the selective correlations among factors at the latent level predicts (*i*) partially separable neurophysiological and genetic mechanisms underlying different forms of endogenous and exogeneous inhibitory control, (*ii*) shared variance among shifting and learning processes, and (*iii*) an intimate relationship about working memory and updating processes. Moreover, the groupings of the six subjects into cognitive phenotypes with niche weaknesses in only a subset of executive subfunctions or overall superior performance across domains provides a starting point to understand the origins of interindividual differences of mental abilities and how they may relate to clinical alterations in mental disorders (*1, 13, 26, 47*).

## Methods and Materials

### Ethics Statement

All animal and experimental procedures complied with the National Institutes of Health Guide for the Care and Use of Laboratory Animals and the Society for Neuroscience Guidelines and Policies and were approved by the institutional review board Vanderbilt University Institutional Animal Care and Use Committee (IACUC) (approval number: M1700198).

### Experimental Procedures

Six rhesus monkeys performed four tasks in 264 experimental sessions on Kiosk touchscreen stations in their home cages (sessions per subject: B: 44; F: 49; I: 44; R: 43; S: 26; W: 58). The Kiosk touchscreen station used the multi-task M-USE software platform to control the visual stimuli, register behavioral responses and trigger water reward as described in detail before(*24, 34*). M-USE (m-use.psy.vanderbilt.edu) is an open source platform based on the Unity video-engine and the four tasks used in this study are part of the default set of cognitive tasks documented and freely available online(*34*). The animals were pair-housed with each pair occupying at least four apartment cages. For each pair of subjects, the Kiosk touchscreen station was mounted as the front panel of one of the apartment cages. The behavioral testing of the animals commenced once they had reached consistent above-chance performance on each of the four tasks and consistently completed all four tasks with the same number of trials per task for each subject (**Fig. 1**, **Suppl. Fig. S1, S2**). Subjects were tested on each weekday at the same time of the day. The same subject of each pair gained access to the Kiosk apartment cage first, and after ∼120 minutes the second subject gained access to the Kiosk. At any time only one of the subjects had access to the touchscreen and the subjects were free to engage with the touchscreen during a 120 min window. All subjects immediately engaged with the touchscreens once the apartment cage with the Kiosk station was opened to them(*24*). The initialization multi-task selection screen showed all four tasks as squared thumbnails on the screen. The task thumbnail of one of the tasks was fully visible and the other three task thumbnails were shown semi-transparent indicating that only one of the four tasks could be started. This ensured the tasks were initiated and performed in a fixed order every session. Subjects started a task by touching its thumbnail. Once a task was completed the screen displayed the multi-task selection screen again with the thumbnail of the next task fully visible and ready to be initiated and all other thumbnails visible semi-transparently and inactive. The thumbnail for each task was a small ∼3cm square image that was generated with the image generation program DALL·E 2 and corresponded to a task-specific theme that was also used as background image for the trials of the task (themes for the four tasks were: skies, sands, clouds, rocks). The unique background contexts were irrelevant to the task and introduced to potentially ease recognition of the specific task.

During task performance subjects received on average 189.42 ± 3.76 mL (38 – 335 mL, n=329) of water per session, dispensed as drops from a water pump through a sipper tube centered in front of the display. The sipper tube extended ∼35 cm from the monitor and ensured that subject maintained a similar distance to the screen throughout task performance. **Movie’s S1-4** show example performance of the subjects, documenting that all subjects sat in front of the monitor, typically with one or two hands on the sipper tube, and reaching towards the screen to initiate trials and chose objects on the screen with a 1-3 finger touch. Consistent touch behavior was trained to the subjects with a separate touch-hold-release task as described before(*34*). The example trials of the videos also show that subjects differed where they placed their hand in between choices/touches, which resulted in a different distance of the hand to the monitor of different subjects. Shorter distances of the hand to the monitor lead to faster reaction times, which is one factor resulting in correlations among reaction times between tasks (**Suppl. Fig. S4**). Because of the subject-specific hand position, we focus our main analysis on accuracy-based metrics.

### Reinforcement procedure

Task performance was based on a positive reinforcement schedule involving collecting secondary symbolic rewards in the form of tokens or slider bar progress and exchanging the secondary rewards for primary water reward once a visually displayed token bar was filled or the slider progressed to the rightmost position. For three tasks, subjects received circular green tokens as symbolic rewards that filled placeholders of a token bar on top of the screen. Once the token bar was filled, it flashed, fluid was dispensed through the sipper tube and the token bar was reset to contain either no or a few tokens. This approach had three purposes. First, the appearance of tokens directly at the touch location on the screen provided immediate visual feedback at the choice location with green tokens for correct and grey tokens for incorrect choices directly above the touched object. The temporal and spatial vicinity of feedback provide maximal salience. Second, the use of tokens instead of immediate fluid rewards allowed analyzing the effects of gaining as well as of losing token rewards on performance, which was used in the Flexible Learning task (*see* below). Third, the use of tokens allowed dispensing water not on every trial but only on trials at which the additional tokens gained for correct choices filled the token bar. This feature facilitated adjusting the number of trials needed for analysis without altering the total fluid amount provided in a given task. Instead of a token bar one task (the Continuous Updating task) used the incrementing of a green slider position to indicate progress towards receiving water rewards. The slider system acted similar to the token bar but provided faster feedback as it did not involve displaying tokens and animating their motion towards the token bar.

With the token- and slider-based reinforcement procedure subjects received actual water reward on average on 463 trials per session (454.5, 640.9, 450.6, 373.7, 320.1, 508.0 for subjects B, F, I, R, S, W). The experiment commenced once all subjects performed at consistent subject-specific performance levels in each of the four tasks and completing the same number of trials within a comparable time window of ∼120 min. While stable performance levels were reached in each of the task (**Fig. 1**, **Suppl. Fig S1**, **S2**) there were subject-specific differences in the overall fluid reward for each task and across sessions. Across all n=329 experimental testing sessions, the Delayed Match-to-Sample task provided on average 21.33 mL fluid reward (21.55, 21.28, 25.84, 24.38, 15.74, 18.31 for subjects B, F, I, R, S, W); the Continuous Updating task provided on average 47.87 mL fluid reward (34.31, 105.14, 44.21, 23.82, 33.41, 45.29 for subjects B, F, I, R, S, W); the Antisaccade task provided on average 83.65 mL fluid reward (41.91, 91.85, 129.26, 121.65, 18.99, 84.58 for subjects B, F, I, R, S, W); and the Flexible Learning task provided on average 36.56 mL fluid reward (29.53, 28.40, 54.71, 44.50, 28.86, 32.95 for subjects B, F, I, R, S, W).

### Task Paradigms and Cognitive Metrics

Four cognitive tasks are administered in a fixed order for all subject in every session: First, the Delayed-Match-to-Sample (DMTS) Working Memory task was performed for 160 trials; second, the Continuous Updating (CU) task was performed for 50 blocks, each with on average 4.47−10.18 trials across subjects which varied dependent on the trial on which subjects committed the first error (*see* below); third, the Antisaccade (AS) task was performed for 207 trials; fourth, the Flexible Learning (FL) task was performed for n = 16 blocks with 25-37 trials each block. For every session, new experimental configurations were generated that ensured randomized feature constellations for objects, randomized object locations, pseudo-randomized sequences of task conditions, etc. The number of trials and blocks of each task were determined prior to the start of the experiment based on three objectives. First, the number of trials/blocks should be sufficiently large to obtain reliable performance metrics for each trial/block within each session. Second, the number of trials/ blocks were set to be as few as needed to minimize the duration of performing the task. Third, the number of trials/ blocks used in the experiment should be identical for each subject to facilitate statistical analysis of inter-individual differences.

All tasks used the same type of 3D-rendered, multidimensional Quaddle 1.0 objects that are generated by matlab (The Mathworks) and python scripts and rendered with the freely available software Blender(*71*). Quaddle 1.0 objects can vary in up to eleven features of the four feature dimensions colors, arm styles, body shapes, and body surface pattern(*71*). The resulting large number of unique feature constellations enabled using new sets of objects for each task and each session. This aspect of the experimental procedures minimizes long-term biases towards specific objects, which is relevant for working memory tasks and the set shifting and learning tasks. An exception to this rule was the Antisaccade task (*see* below) that used a pre-defined set of four uniform, grey-colored Quaddle 1.0 objects with different body shapes (hourglass, oval, cylindrical, cubical)(*71*).

### Accuracy-based Antisaccade task

The accuracy-based Antisaccade (AS) task measured how accurate subjects process a briefly presented object opposite to a cued location by asking subjects to recognize the presented object in an array of 4 objects. Example performance is shown in **Movie S1**. Each trial randomly used one of four pre-defined objects - with body shapes like an hourglass, oval, cylindrical, or cubical - as the target and the remaining objects of the object set as distractors. The task involved choosing an object by touching it on a touchscreen and not by eye movements, but was named antisaccade following conventions in the field(*6*). A trial started with a centrally presented start button and a centrally presented alert cue. After an initial 0.5 s delay a peripherally presented cue stimulus (asterisk) was shown for 0.15 s cue followed 0.05 later by a grey-colored target object that could be presented at the cue location (prosaccade trial) or opposite to the cued location (antisaccade trial). Following a 0.5 s delay four objects were presented at random locations of a virtual circle equidistant to other objects and to the center of the display. Overall accuracy and reaction times in the antisaccade condition was calculated as *AS_Accuracy* and *AS_ProcessingSpeed* (**Fig 1B**,**C**). The relative decrease in accuracy on the anti-versus pro-saccade condition was calculated as the index (Antisaccade - Prosaccade / Antisaccade + Prosaccade) to evaluate a normalized difference score in metrics named *AS_Accuracy_ProAnti_Index(i)* and *AS_RT_ProAnti_Index(i)*. The suffix ‘(i)’ signifies - for this and all other metrics - that the metric values were inverted so that more positive values reflect better performance (i.e. less interference). The response display could show the target object in the same, congruent hemifield as the target location, or in the opposite, incongruent hemifield, which was used to quantify the spatial congruency Simon effect as (Incongruent - Congruent / Incongruent + Congruent), captured by the metric (*AS_Accuracy_SimonCong_Index(i)*, *AS_RT_SimonCong_Index(i)*). We also used the average accuracy score across anti- and prosaccade conditions to avoid a bias toward only considering antisaccade trials as is done in some human studies (metric: *AS_PS_Accuracy*).

For the AS task as well as the other tasks we leverage both, overall accuracy metrics as well as normalized difference metrics. The use of difference score metrics is supported by a recent meta-analysis of human attentional control studies that concluded understanding attentional control functions can utilize either difference scores or avoid difference scores with rather minimal differences in cognitive factors inferred from models that use exclusively difference scores or avoid them(*7*). The rationale for using the AS task is its long history as a behavioral task isolating the ability to exert voluntary control over reflexive orienting (*72*) and the more recent finding in human studies on cognitive factors that the AS task consistently showed highest intra-subject consistency and higher loading on a latent attention (inhibitory) control factor than other conflict task(*6, 42*). In addition, inter-individual variations of AS task performance gain importance because they are indicative of multiple neuropsychiatric conditions that involve altered recruitment of the prefrontal cortex – striatum network(*73*).

### Continuous Updating Task

The Continuous Updating (CU) task presented subjects blocks with trials in a block displaying one additional object every new trial. The first trials showed two objects, the third trial showed three objects, etc. Subjects had to choose an object they did not chose in prior trials of that block, i.e. they followed the non-matching rule ‘Don’t pick an object twice’. Once subjects committed an error by choosing an object they had chosen in a prior trial the block ended and a new block started. Example performance is shown in **Movie S2**. Each session presented 50 CU blocks. Each block began with a trial presenting two objects at random locations and subjects could choose either of those objects to get reward. The next trial presented the same two objects plus an additional object and subjects were rewarded when they chose the object not previously chosen. The third trial presented at random locations all prior presented objects plus an additional object and subjects were rewarded by choosing an object that was not chosen in prior trials. This procedure continued for up to 20 trials, i.e. up to 21 objects displayed on the screen. When a subject incorrectly chose an object they had previously chosen, the block ended and the trial number when this error occurred was registered. The trial number at which an error was committed in 50% or in 75% of blocks served as a metric quantifying working memory updating span (metrics: *CU_Accuracy_AvgTrialToError*, *CU_Accuracy_TrialAt75Acc*) (**Fig. 1E**). Subjects’ performance across 50 blocks in a session showed a characteristic trend of increasing likelihood to commit an error with increasing number of stimuli. A regression line fitted to this effect quantified how much accuracy declined with growing numbers of objects in working memory (i.e. growing numbers of objects in the display that were chosen in prior trials) (metrics: *CU_Accuracy_Slope*). Reaction times increased with increasing number of objects on the screen, which we quantified with a regression slope indexing a working memory set size effect, and with a regression intercept indexing the overall processing speed (metrics: *CU_ProcessingSpeed_Slope*, *CU_ProcessingSpeed_Intercept(i))* (**Fig. S1H-J**). When subjects erroneously chose an object they had chosen in prior trials, this erroneously chosen object might have been an object they chose in the most recent past (e.g. in the last trial t_-1_), or it might have been an object they had chosen several trials prior to the trial with the error (e.g. trials t_-2_, t_-3_, … t_-n_). We quantified the probability that the erroneously chosen object was t_-1_, t_-2_, … t_-n_ away. A positive regression slope over this probability indexes that errors were more likely committed for objects more recently chosen, i.e. that errors indicated confusion of the most recently updated objects in working memory. We captured this with the performance metric *CU_nBackSlope(i)*. This analysis showed that subjects most likely erroneously chose a recently chosen object (**Fig. 1F**). The rationale for using the CU task is based on insights of cognitive flexibility models that distinguishes contributions of continuous monitoring and of updating of working memory contents(*74*). In addition, the updating of working memory content has been found to be separated from processes linked to response inhibition (Chase et al., 2008; Friedman et al., 2006; Miyake et al., 2000). The CU task is similar to the Delayed Recognition Span task (Moore et al., 2005) and to self-ordered search tasks (Axelsson et al., 2021; Champod & Petrides, 2007; Walker et al., 2009) used in prior NHP studies. Performance of the CU task has clinical significance as deficits in self-ordered selection tasks are evident in humans diagnosed with attention-deficit-hyperactivity disorder (ADHD) (Dowson et al., 2004; Fried et al., 2015) and schizophrenia (Badcock et al., 2005; Pantelis et al., 1999), including first degree relatives and monozygotic twins of schizophrenic patients (Pirkola et al., 2005; Wood et al., 2003) even when they have no altered cognitive functioning (Joyce et al., 2005).

### Flexible Learning Task

The Flexible Learning (FL) task combines an attentional set shifting task, requiring switching between familiar feature-rules, with a feature-based attentional learning task, requiring learning new feature-reward rules (for details *see*(*75*)). The task is described in detail in (*75*) and example performance is shown in **Movie S3**. In each trial, three Quaddle objects are displayed on a virtual circle equidistant from each other and the monitor center that vary in features of two feature dimensions (e.g. colors and body shape). One of these features is a designated target and associated with reward. Subjects had to choose an object each trial and if the chosen object contains the target feature subjects receive either +2 or +4 tokens (conditions: *Gains +2, +4*), and otherwise they lose 1 or 3 tokens for choosing objects with incorrect features (conditions: *Loss -1, -4*). The rewarded target feature stayed constant for blocks of 25-37 trials. Subjects learned by trial-and-error which object feature is the target. In each session, there were 16 different blocks with pseudo-randomized block-transitions that either associated a novel feature with reward that was not shown in the previous block (block transition condition: *New*), or re-assigned one feature that was a non-rewarded distractor feature in the previous block to become the target feature in the new block (block transition condition: *Same*). Independent of the New/Same transition rules the task also varied whether the newly rewarded target feature was of the same feature dimension as the target of the previous block (block transition condition: *Intradimensional ID*) or was a feature from a different feature dimension as the target of the previous block (block transition condition: *Extradimensional ED*). An ID example is a change from an oblong shaped target to a pyramidal shaped target, and an ED example is a change from an oblong shape to a checkerboard body pattern as target feature. In addition, each block was pseudo-randomly assigned a gain for correct choices (conditions: Gains +2, +4), and a loss for erroneous (non-target) choices (conditions: Loss -1, -3). For example, in a block with Gain +2/Loss-3 subjects would receive 2 reward tokens when choosing the target and loosing 3 already attained reward tokens when choosing a non-target object. A block started with a token bar that had 8 placeholders for tokens and 3 tokens as starting asset. Once subjects filled the 8 token placeholders, they received reward and the token bar was reset to contain 3 starting tokens. We analyzed how performance within blocks varied according to those conditions with three different performance metrics. First, we quantified how fast subjects learned the rewarded target by quantifying the number of trials needed to reach criterion performance, which we defined as the first trial within a block in which a forward-looking 10-trial window had 70% accuracy (metric: *FL_LearningSpeed(i)*). We then quantified how the trials-to- criterion (T2C) changed for the different block-transition conditions by calculating a normalized index *(Same_T2C_ - New_T2C_ / Same_T2C_ + New_T2C_)* for the metric *FL_LearningSpeed_Novelty_Index(i)*; the index *(ED_T2C_ - ID_T2C_ / ED_T2C_ + ID_T2C_)* for the metric *FL_LearningSpeed_Dimension_Index(i)*; the index *(Gain2_T2C_ – Gain4_T2C_ / Gain2_T2C_ + Gain4_T2C_)* for the metric *FL_LearningSpeed_Gain_Index(i)*; the index *(Loss1_T2C_ – Loss3_T2C_ / Loss1_T2C_ + Loss3_T2C_)* for the metric *FL_LearningSpeed_Loss_Index(i).* Second, we calculated the overall proportion of rewarded choices (accuracy) over trials starting from the first trials at which criterion performance was reached to the last trial in a block. This plateau accuracy is captured by the metric *FL_Accuracy* (and for the overall plateau reaction time: *FL_ProcessingSpeed(i)*). We then calculated the normalized difference scores for accuracy across different block transition conditions in the same way as for the learning speed to arrive at the metrics *FL_Accuracy_Dimension_Index(i)*, *FL_Accuracy_Novelty_Index(i)*, *FL_Accuracy_Gain_Index(i)*, and *FL_Accuracy_Loss_Index(i)*. Third, we extended the metrics on learning and plateau accuracy by a trial-level metric that quantified the probability of subjects to make two or more consecutive errors (E_n_) after a correct (C) choice. This CE_n_ analysis evaluated perseverative errors with the metric *FL_CEn_Perservation(i)* and is an important metric used to assess prefrontal cortex function(*76*). To measure how CE_n_‘s change in different block transition conditions we calculated the same normalized difference index as for trials-to-criterion and Accuracy also for the CE_n_ measure in the metrics *FL_CEn_Perservation_Dimension_Index(i)*, *FL_CEn_Perservation_Novelty_Index(i)*, *FL_CEn_Perservation_Loss_Index(i)*, *FL_CEn_Perservation_Gain_Index(i)*. The rationale for using the FL task is based on studies distinguishing a latent shifting factor (*see* Discussion) and on its similarity to the previously used Wisconsin Card Sorting Task, the Category Set Shifting Task and to classical Extra-/Intra-dimensional shifting tasks (*77, 78*) used in prior NHP studies.

### Delayed Match-To-Sample Task

The Delayed Match-To-Sample task (DMTS) tests how well subjects sustain visual objects in short-term memory over delay. Example performance is shown in **Movie S4**. The task presented a sample Quaddle object for 0.5s, followed by a delay of 0.3, 1.3, 2.3, 3.8, or 5.1s and a response display that contained the sample object and other distracting Quaddle objects at random locations. The subject had to choose the response display object that matched the sample object. Performance accuracy and reaction-times (RT) were analyzed with metrics estimating short-term memory effects and the effects of different types of interference from distraction (*see* **Fig. 1K**,**L****, SG-E**): (**1**) The task manipulated the delay to measure how accuracy/RT declined with the duration passed since the sample disappeared from the screen, measured as the slope of a regression over the delays, quantified with the metric named *DMTS_Accuracy_Slope*, *DMTS_ProcessingSpeed(i)*. The suffix (*i*) signifies - here and for all other metrics - that the obtained slope value was inverted so that more positive values reflect better performance (here: that shallower slopes reflect less decline in performance over delays, i.e. better short term memory maintenance). (**2**) The task used Quaddle objects as targets and distractors that varied in features of one or two different feature dimensions (e.g. varying in color and body shape or pattern of the object body). The task contained two conditions in which targets and distractors shared features of either one or two dimensions which reflect low or high target-distractor similarity (TDS). We computed how TDS similarity affected performance with an index for accuracy/RT (High_TDS_ - Low_TDS_ / High_TDS_ + Low_TDS_) ranging from +1 to -1. We inverted this TDS index so that higher values correspond to being less affected by perceptual interference (metrics: *DMTS_Accuracy_TDS_Index(i)*, *DMTS_RT_TDS_Index(i)*). (**3**) During the delay we presented in one third of the trials a distractor object for 0.5 s at the location at which the sample was presented (*P*ost-*S*ample-*D*istractor: *PSD*) to evaluate how accuracy/RT declined in trials with versus without interference from a distractor, which we indexed as (With_Distractor_ – Without_Distractor_ / With_Distractor_ + Without_Distractor_) (metrics: *DMTS_Accuracy_PSD_Index(i)*, *DMTS_RT_PSD_Index(i)*). (**4**) Subjects had to find and chose the sample object in the response display. We populated the response display with the sample target object and either with two or three distractor objects to test whether increased interference from an additional third distractor object affected performance. We calculated this interference effects with the index (Three_NumDistractor_ - Two_NumDistractor_ / Three_NumDistractor_ + Two_NumDistractor_) for accuracy and RTs (metrics: *DMTS_Accuracy_NumDist_Index(i)*, *DMTS_RT_ NumDist _Index(i)*). (**5**) The DMTS task also provided performance metrics for the overall accuracy (*DMTS_Accuracy*) and overall processing speed indexed as the intercept of the regression of reaction times over delay durations (*DMTS_ProcessingSpeed_Intercept(i)*).

### Evaluating normalized cognitive profiles

The four tasks were analyzed according to 33 behavioral metrics (**Fig. 2B**). For all metrics a more positive, larger value indicated improved performance/less interference/faster responses, which was ensured by inverting all metrics in which the lower score would have indicate better performance. We added the suffix ‘*(i)*’ for all metrics where the raw metrics were inverted to indicate superior performance with more positive values (e.g. a lower number of trials-to-criterion indexing faster learning in the Flexible Learning task; or a lower difference in accuracy between anti- and prosaccade trials indexing less decline in performance in the antisaccade condition). To compare performance across individuals and metrics, we first standardized the raw scores for each variable into z-scores across the entire dataset. We then calculated the mean z-score and standard error of the mean (SEM) for each subject by averaging these standardized scores across sessions (**Fig. 2D**).

### Assessment of reliability of behavioral metrics

We assessed the consistency of performance scores by calculating the Spearman-Brown split-half reliability for each metric (**Fig. 2C**). We implemented a bootstrapping procedure with 500 iterations for each metric. In each iteration, every subject’s set of completed sessions was randomly partitioned into two equal halves. The mean performance score for each half was calculated, and a Pearson correlation was computed across subjects between the scores from the two halves. This correlation (r) was then adjusted using the Spearman-Brown prophecy formula to estimate the reliability of the full measure. If an iteration produced a negative correlation, reliability was set to 0 for that split. The final reliability score for each metric is the mean across the 500 iterations. Metrics with a mean reliability score below a threshold of 0.5 were excluded from further analysis.

### Unsupervised discovery of cognitive phenotypes

To investigate whether subject-wise differences in task performance formed subject specific cognitive phenotypes, we used an unsupervised discovery approach that combined dimensionality reduction with clustering (**Fig. 3**). All analyses were performed in Python using the scikit-learn and umap-learn libraries. First, to ensure data integrity, performance metrics were standardized to zero mean and unit variance to prevent feature scaling biases. We then projected the high-dimensional feature space into a two-dimensional embedding using Uniform Manifold Approximation and Projection (UMAP). The UMAP algorithm was configured with 15 nearest neighbors and a minimum distance of 0.1 to preserve both local and global data structures.

Following dimensionality reduction, k-Means clustering was applied to the 2D UMAP coordinates. To determine the optimal number of clusters, we utilized the Elbow Method, evaluating the Sum of Squared Errors (SSE) across a range of cluster numbers (k = 1 to 10) (**Fig. 3B**). The optimal k was identified as the point of maximum curvature using the Knee Locator algorithm. The final K-Means clustering was run with the determined k, using k-means++ initialization and 10 random restarts to ensure the stability and robustness of the cluster assignments. We assigned each experimental session a cluster ID to allow visualization and subsequently quantified the proportion of sessions belonging to each cluster for every subject (**Fig. 3C**). To characterize the functional profile of these phenotypes, we calculated the mean performance for overall task accuracy metrics within each cluster. To facilitate cross-task comparison, these cluster means were standardized (Z-scored) relative to the distribution of means across all identified clusters, with error bars representing the standard error (**Fig. 3D**).

### Disentangling intra- and inter- subject variability and performance

Given our repeated-measures, small-N design, a critical analytical step was to disentangle two primary sources of performance variation: inter-subject differences in cognitive abilities, and intra-subject fluctuations in performance from session to session. To isolate the within-subject relationships, we computed partial Pearson correlations for all pairs of reliable behavioral metrics (**Fig. 4**, **S6**). This was achieved using a linear mixed-effects (LME) model framework for each pair of metrics.

In a complementary repeated-measures analysis, we quantified the subject-level contribution to the fourteen-metrics contributing to the exploratory factor analysis results (*see* below). For each metric and each session-level EFA factor score, we estimated a one-way random-intercept variance decomposition with subject identity as the grouping factor. Intraclass correlations were calculated as the between-subject variance divided by the sum of between-subject and within-subject/session variance. Details of the variance decomposition are described in **Supplementary Methods:** *Variance decomposition for latent factor analysis*, and in **Suppl. Fig. S6**.

### Exploratory factor analysis to quantify latent factors underlying Executive Functions

To identify the underlying structure of cognitive abilities, we performed a latent variable analysis using a three-stage approach: (1) data suitability testing, (2) Exploratory Factor Analysis (EFA) to model latent constructs, and (3) hierarchical clustering to visualize metric similarities. All analyses were conducted in Python using the factor_analyzer and scikit-learn libraries (https://factor-analyzer.readthedocs.io/en/latest/index.html and https://scikit-learn.org/stable/). To ensure the dataset was suitable for factor analysis, we conducted an iterative Kaiser-Meyer-Olkin (KMO) test. Starting with all reliable metrics, we calculated the Measure of Sampling Adequacy (MSA) for each variable. In each iteration, the variable with the lowest MSA was removed if it fell below the threshold of 0.5. This process was repeated until all remaining variables met the minimum MSA criterion, ensuring a robust dataset for structural analysis. The EFA was fitted to the fourteen pooled standardized metrics using a four-factor varimax-rotated solution (**Fig. 5A**).

We conducted a formal EFA on the KMO-filtered dataset to identify latent cognitive factors(*40, 79*). The number of factors to extract was determined objectively using the Kaiser criterion, retaining factors with eigenvalues greater than 1 as derived from an initial Principal Component Analysis (PCA). Factor extraction was performed using the principal factor method. To account for the theoretical likelihood that cognitive factors are interrelated rather than independent, we applied an oblique Promax rotation. Factor loadings were visualized to interpret the semantic meaning of each latent dimension (**Fig. 5A**). Correlations among latent factors were calculated as the off-diagonal elements of the resulting factor correlation matrix, representing the cosine of the angle between the oblique factor axes (**Fig. 5B**). To validate the latent factorial structure we performed hierarchical clustering on the loadings of the first two principal components (**Suppl. Fig. S5**). The resulting relationships were visualized as a dendrogram, which depicts the similarity structure and nested clusters of the performance metrics based on their Euclidean distance in the reduced PCA space.

### Additional testing for a common general factor

To test whether our dataset supports the existence of a general common factor in addition to the specialized latent factors that were obtained from the exploratory factor analysis we used multiple approaches (see **Supplementary Methods**). An exploratory structural equation model (ESEM) was used to test how cross-loadings of latent factors would emerge; Confirmatory factor analyses (CFA) and bifactor analyses (BF) were used to test how assuming a fixed shared resource could explain variance across metric; Higher-order general-factor analysis was used to test whether a common general factor may be expressed though specialized factors in a hierarchical variance structure that. These approaches are summarized in **Suppl. Methods**. These approaches provided converging evidence against a reliable common factor in our dataset (**Suppl. Fig. S7**; **Suppl. Table S1**, **S4, S6**). We also extended the analysis by considering reaction time measures in addition to the fourteen reliable accuracy-based metrics, which likewise did not suggest a common factor (**Suppl. Results**).

## Financial Disclosures

The authors declare no competing financial interests.

## Acknowledgements

We thank Gordon Logan for helpful comments on the manuscript. This work was supported by the National Institute of Mental Health of the National Institutes of Health under Award Number R01MH129641 (TW). The content is solely the responsibility of the authors and does not necessarily represent the official views of the National Institutes of Health.

## Data, Materials, and Software Availability

The data and analysis code supporting this study will be available in a public repository on Zenodo upon publication. All other data are included in the article and/or supporting information.

## Author Contributions

A.N. and X.W. performed research; X.W., L.M. analyzed data; X.W., A.N., T.W. edited the paper; T.W. wrote the paper; T.W. conceived and supervised the project

## Notes

### Competing Interest Statement

The authors have declared no competing interest.

### Summary of Updates

- Introducing new confirmatory factor analyses testing explicitly for a common factor versus multiple latent factors. - New quantification of the influence of between- and within- subject variance to account for the latent factor structure of the exploratory factor analysis. - Clarifications in the introduction, results, discussion and methods sections.

