## Supplementary Material for "Inhibitory Control, Shifting, and Working Memory Updating Domains form Cognitive Phenotypes in Non-human Primates"

**Content:**

**Supplemental Results**

**Supplemental Figures S1-S9**

**Supplemental Table**

**Supplemental Movies S1-S4**

### Supplemental Results

#### Four-factor structure is influenced by within- and between-subject variance

The exploratory factor analysis provided a four-factor structure that we interpret as originating mostly from between-subject variance. However, the factor analysis result may not only be influenced by differences between subjects but also by within-subject variance across sessions. To quantify these effects we performed a variance decomposition analysis of the fourteen accuracy-based metrics. We found that most metric variance was within-subject/session variance rather than between-subject variance (median metric intraclass correlation (ICC) = 0.174). The same result pattern was evident for the exploratory factor analysis (EFA) factor scores, for which the median within-subject/session variance share was 66.8% (**Suppl. Fig. S8A-C**). However, the 4-factorial loading structure was preserved when the stable subject means were removed: The metrics correlation matrix was similar when performance scores were pooled across subjects (pooled matrix) compared with the within-subject centered matrix (upper-triangle r = 0.873). The pooled four-factor loading pattern showed high best-matched Tucker congruence with the within-subject-centered solution (mean |Tucker φ| = 0.91) (**Suppl. Fig. S8D-H**). Thus, the four-factor structure was not solely explained by stable between-subject differences, although the between-subject structure remains descriptive given the sample of six subjects.

We also tested whether adding reaction-time and processing-speed metrics would reveal a dominant common factor. In a processing speed-inclusive 24-metric EFA, factor-retention criteria did not converge on a single solution: MAP favored three factors, the scree elbow favored five factors, parallel analysis favored six factors, and the Kaiser criterion favored eight factors (**Suppl. Fig. S9A,B**). We therefore treated the five-specific-factor bifactor model as a sensitivity analysis. Compared with the fourteen-metric model, adding speed-related metrics increased the apparent general-factor contribution, but the general factor remained below common interpretability thresholds (14 metrics: ECV = 0.245, $\omega_h$ = 0.304; 24 metrics: ECV = 0.352, $\omega_h$ = 0.494) (**Suppl. Fig. S9C-E**). Moreover, ProcessingSpeed metrics accounted for 73.6% of the summed squared G loadings in the 24-metric model, while RT contrast metrics accounted for 3.2% (**Suppl. Fig. S9F**). This suggests that the speed-inclusive general factor primarily reflects shared processing-speed variance rather than a broad common executive-function factor.

### Supplemental Figures S1-S8

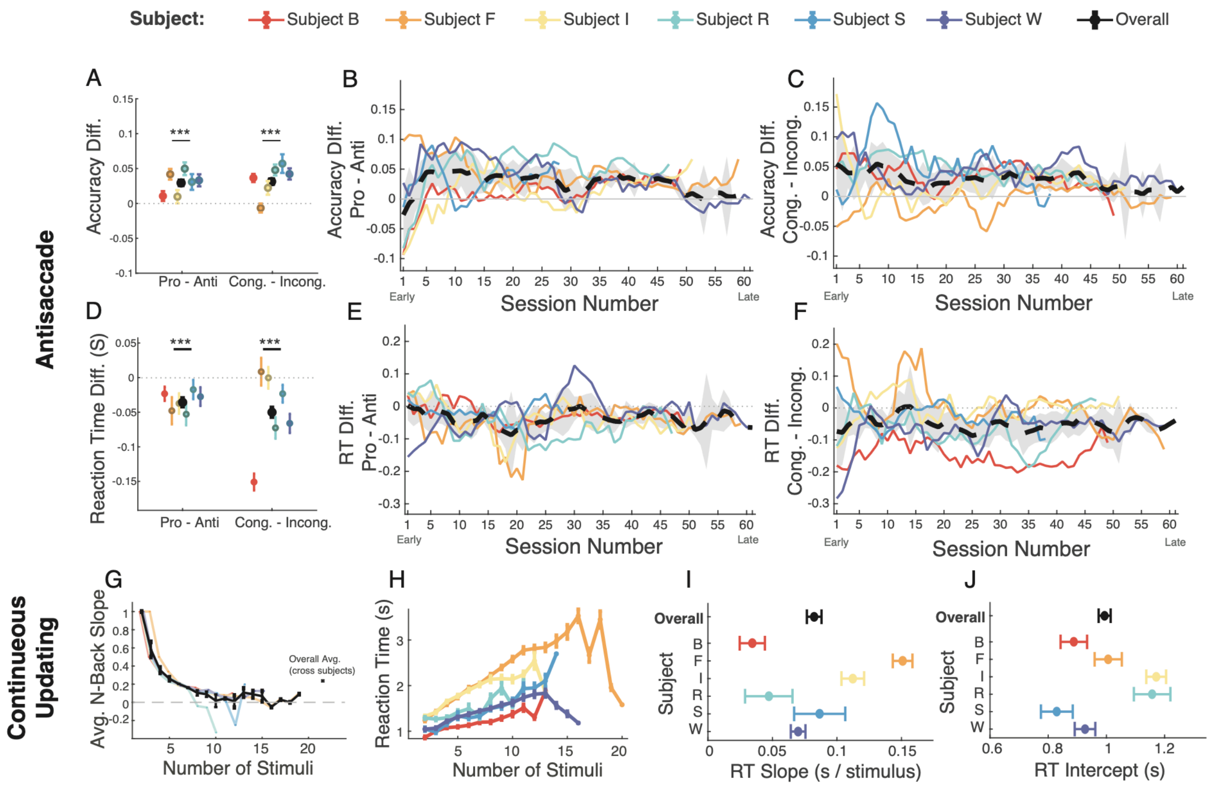


**Suppl. Fig. S1. Extended behavioral performance metrics for the Antisaccade and Continuous Updating task.** (**A**) Mean accuracy difference for Antisaccade Cost (Pro - Anti) and Simon Effect Cost (Cong. - Incong.). Asterisks indicate the group mean (black dot, ± SE) is significantly different from zero (one-sample t-test, $p < 0.001$). (**B**) Antisaccade Cost (Accuracy Diff.) plotted over session number. (**C**) Simon Effect Cost (Accuracy Diff.) plotted over session number. (**D**) Mean reaction time (RT) difference for Antisaccade Cost (Pro - Anti) and Simon Congruency Effect Cost (Cong. - Incong.). Asterisks indicate the group mean (black dot, ± SE) is significantly different from zero (one-sample t-test, $p < 0.001$). (**E**) Antisaccade Cost (RT Diff.) plotted over session number. (**F**) Simon Congruency Effect Cost (RT Diff.) plotted over session number. In (B, C, E, F), colored lines represent individual subjects, the black dashed line is the group average, and the shaded gray area is the SE. (**G**) Average reaction times (RT) of Continuous Updating Task as a function of number of the stimuli displayed. (**H**) Mean slope (± SE) of the linear fit to RT vs. number of stimuli (from G) for each subject and the group overall (black). (**I**) Mean intercept (± SE) of the linear fit to RT vs. number of stimuli displayed (from G) for each subject (colors) and the group (black).

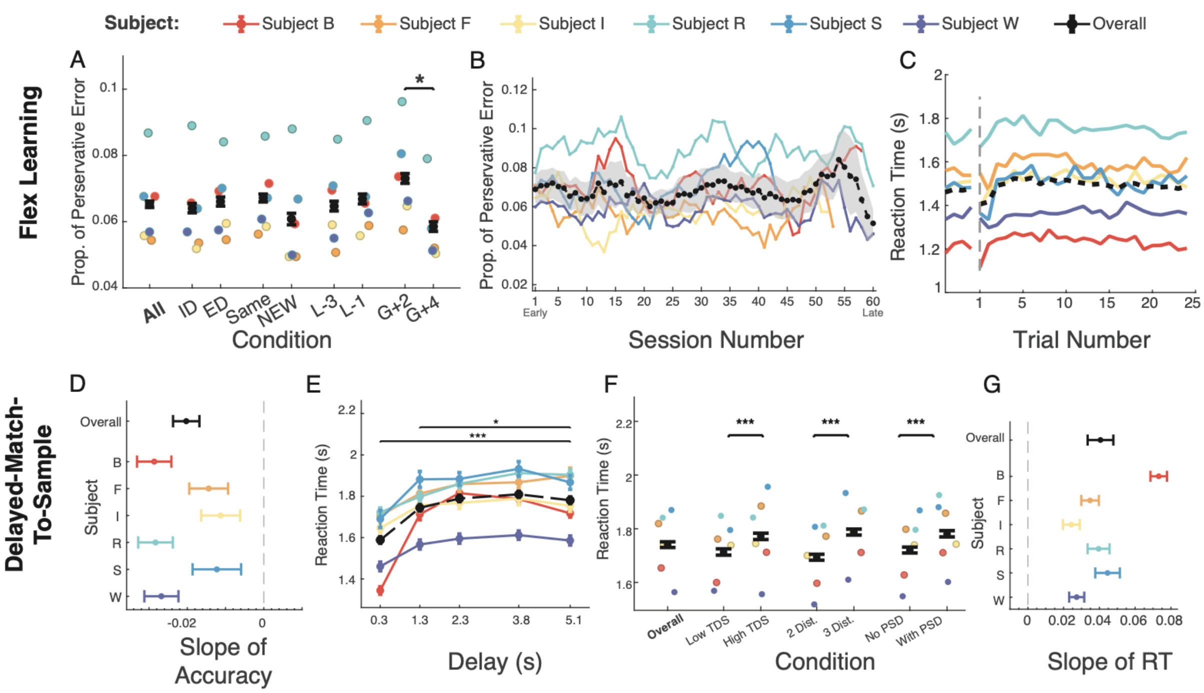


**Suppl. Fig. S2. Extended behavioral performance metrics for the Flexible Learning task and the Delayed Match-to-Sample task** (**A**) Proportion of perseverative errors (defined as errors following correct choices, out of all errors) across different block conditions of the Flexible Learning task. The asterisk indicates a significant pairwise comparison. (**B**) Proportion of perseverative errors plotted over session number. The black dashed line is the group average, and the shaded gray area is the SE. (**C**) Average reaction time (RT) as a function of trial number within a block. Colored lines represent individual subjects, and the black dashed line is the group average. (**D**) Based on performance of the Delayed Match-To-Sample task the plot shows the mean slope (± SE) of the linear fit to accuracy vs. delay duration (from **Fig. 1L**) for each subject and the group overall (black). (**E**) Average reaction time (RT) as a function of delay duration. Asterisks indicate significant pairwise comparisons. (**F**) Average RT across different interference conditions (TDS: Target-Distractor Similarity; PSD: Post-Sample Distractor). Asterisks' indicate significant pairwise comparisons. (**G**) Mean slope (± SE) of the linear fit to RT vs. delay duration (from H) for each subject and the group overall (black).

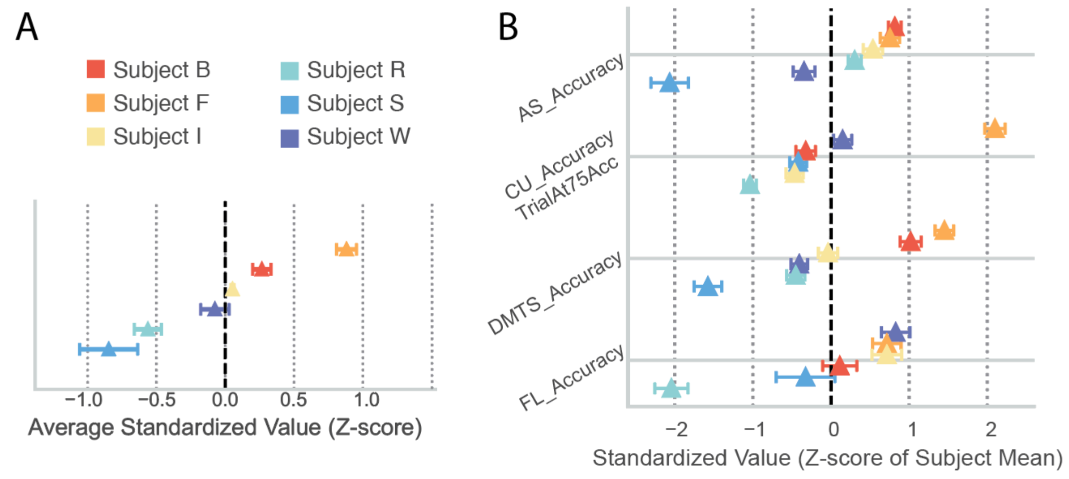


**Suppl. Fig. S3. Normalized performance scores across metrics and each main task metric.** (A) Average z-standardized performance across all reliable performance scores for each subject. (B) Z-standardized performance (mean Z-score ± SE) for the main performance metric of each of the four tasks for each subject.

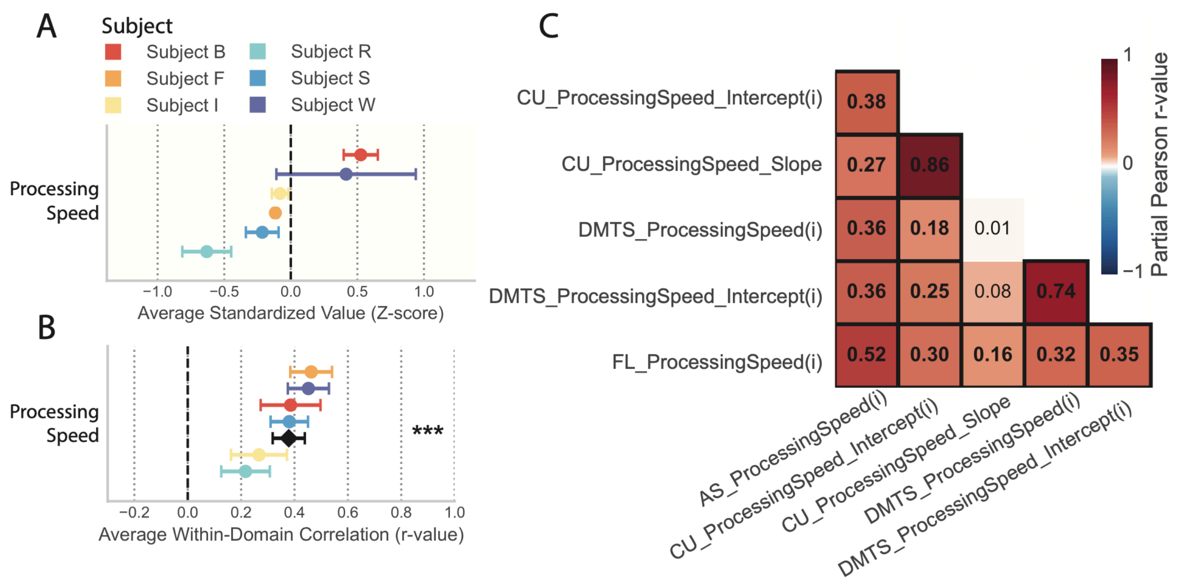


**Suppl. Fig. S4. Normalized reaction times and correlation of reaction time metrics.** (**A**) Average z-normalized (±SE) reaction times across all metrics indexing processing speed (reaction time). (**B**) Average correlation across all metrics indexing processing speed. Individual subjects are shown in colors and the group average in black. (***) indicates $p<0.001$ difference from zero. (**C**) Partial Pearson correlations of the main reaction time metrics. Significant correlations ($p < 0.05$, FDR corrected) have thick black border. A caveat interpreting the apparent correlations are subject differences of how distant the hand was positioned from the display prior to the making a choice, introducing inter-subject variance (see Methods).

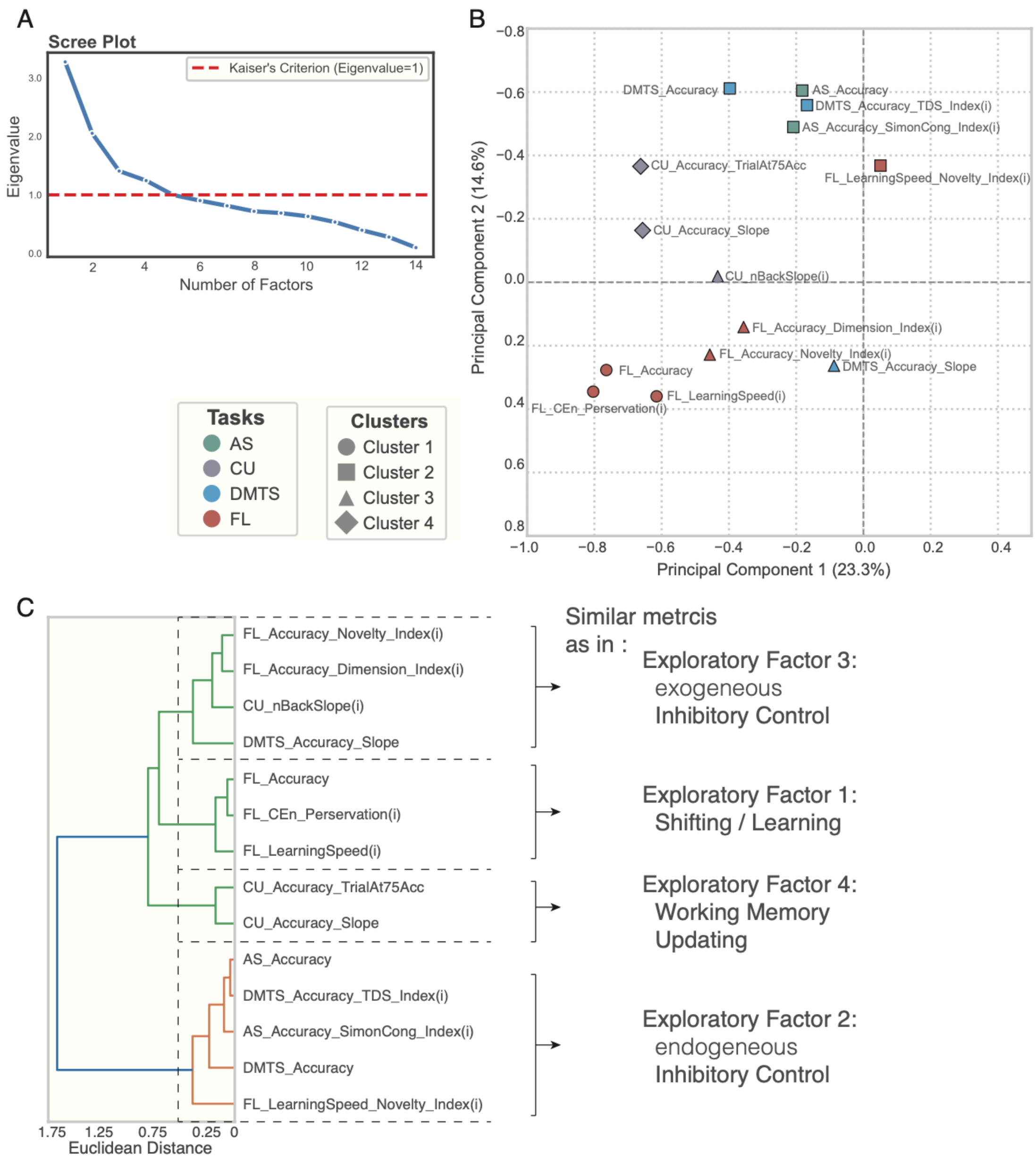


**Suppl. Fig. S5. Number of Latent Factors, Principal Component Analysis (PCA) and Hierarchical Clustering of KMO-Filtered Metrics**. (**A**) The Kaiser's criterion threshold of an Eigenvalue =1 suggests that our dataset is well described with four separable latent factors. (**B**) Loading of the first two principal components of the behavioral metrics. Point color indicates the task, and point shape indicates the cluster assignment (k=4, determined by the elbow method) from K-Means clustering applied to these 2D coordinates. (**C**) Hierarchical clustering dendrogram (Ward's method). The clustering is based on the Euclidean distance between the 2D PCA loading coordinates of the metrics shown in *B*. A dashed vertical line at a distance of 0.5 suggests four separable cluster groups whose metrics are similar to the groupings of metrics in the four latent factors of the exploratory factors as described on the right side of the plot.

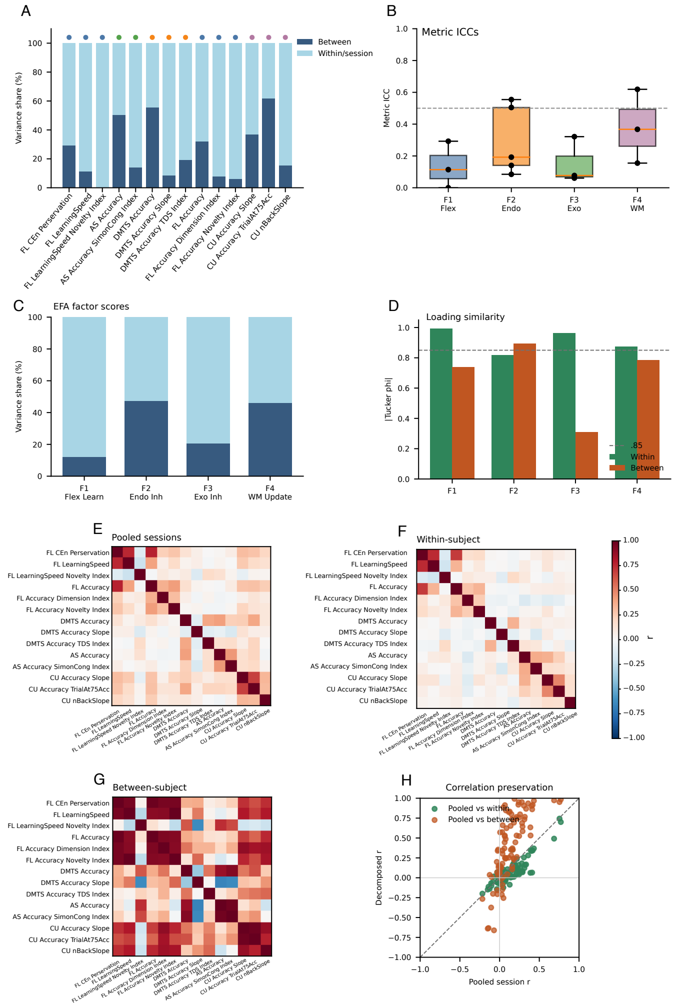


Figure S8

**Suppl. Fig. S6. Within- and between-subject variance decomposition of the 14-metric panel.** (**A**) Variance decomposition for each observed metric, showing the estimated percentage of variance attributable to between-subject differences and within-subject/session variation. (**B**) Metric-level intraclass correlations grouped by the four latent exploratory factors (see **Fig. 5**). (**C**) Variance decomposition of session-level EFA factor scores from the pooled four-factor solution. (**D**) Loading similarity to the pooled four-factor EFA, quantified as best-matched absolute Tucker congruence for within-subject-centered and subject-mean solutions. (**E**) Pooled session correlation matrix, calculated across all available session-level observations after metric standardization, without separating stable subject means from session-to-session variation. This matrix therefore combines both between-subject covariance and within-subject/session covariance. (**F**) Within-subject-centered correlation matrix, calculated after subtracting each subject's own mean from each metric. This removes stable differences among animals and isolates whether metrics covary across sessions within the same subject. (**G**) Between-subject mean correlation matrix, calculated from each subject's mean profile across sessions. This matrix captures whether animals that are high on one metric also tend to be high on another metric but is descriptive because it is based on six subject-level observations. (**H**) Upper-triangle correlation preservation, comparing pooled metric-pair correlations with corresponding within-subject-centered and between-subject-mean correlations. These analyses show that the pooled factor structure was not explained solely by stable differences among subjects, while between-subject inference remains descriptive because the sample included six subjects.

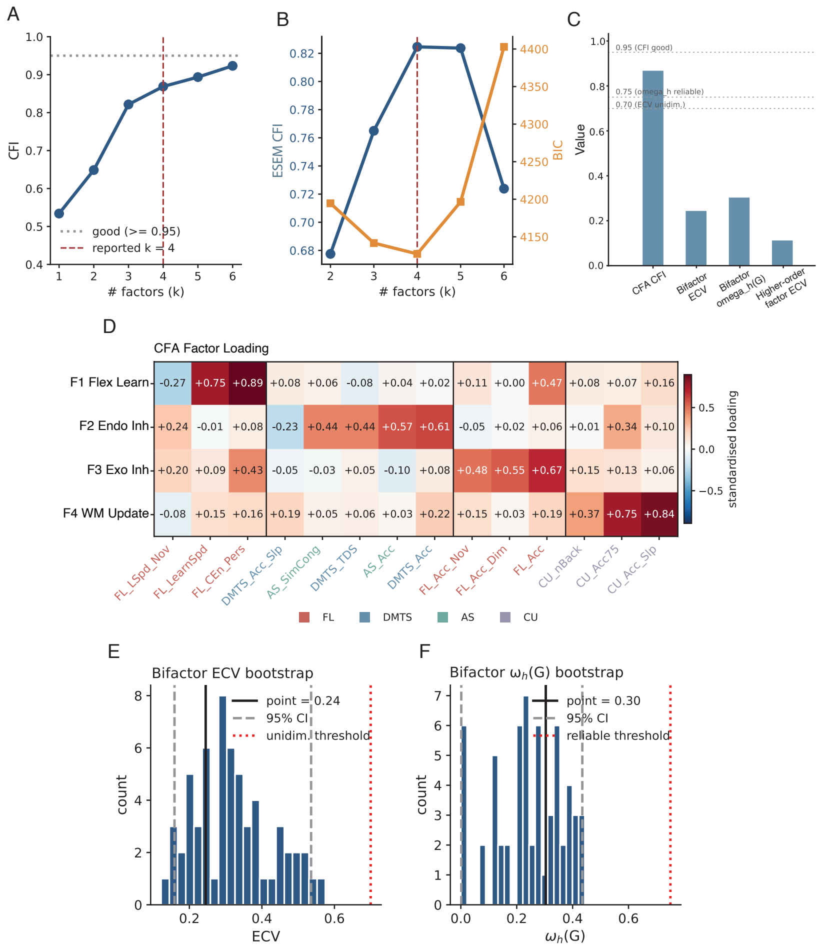


Figure S7

**Suppl. Fig. S7. Convergent support for a four-factor solution and limited support for a single common (general) factor.** (**A**) Confirmatory factor analysis (CFA) model fit across one- to six-factor solutions based on the 14-metrics that we used for the exploratory factor analysis (**Fig 5**). The four-factor CFA model was better than the three-factor solution, but with overall moderate goodness of fit. (**B**) ESEM comparison of models with varying number of factors showed that the Bayesian Information criterion (BIC) was lowest (i.e. best) and ESEM CFI was highest (i.e. best) for the 4-factor solution. (**C**) Summary indices comparing four-factor CFA fit with bifactor and higher-order estimates of a general factor. Bifactor explained common variance (ECV), bifactor omega hierarchical, and higher-order ECV were below conventional thresholds for interpreting a strong general factor. (**D**) Standardized factor-loading matrix for the 14-metric four-factor solution, with metrics ordered by manuscript factor assignment and task family. (**E**,**F**) Subject-stratified bootstrap distributions for bifactor ECV and omega hierarchical of the general factor. Bootstrap estimates remained below common reliability thresholds, indicating that the data support four correlated cognitive factors more strongly than a stable unitary executive-function factor.

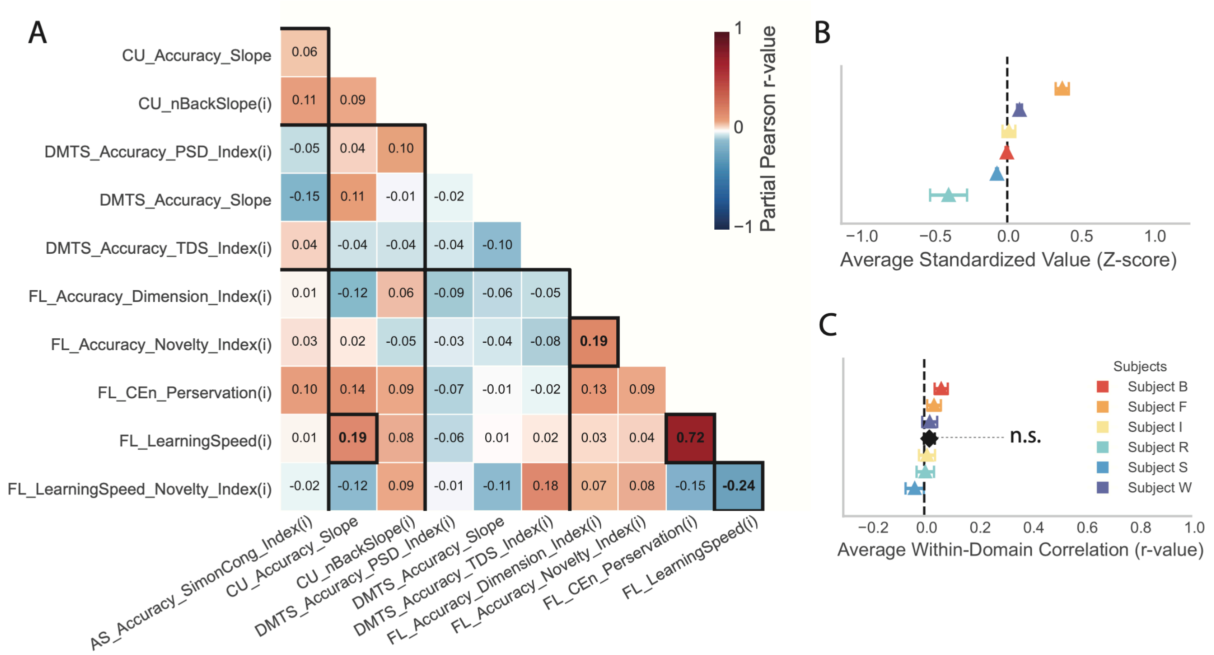


**Suppl. Fig. S8. Correlations of performance scores of metrics that indexes inhibitory control of interference from different tasks.** (**A**) Partial correlations of metrics indexing inhibitory control. Black outlined cells indicate statistical significance ($p<0.05$, FDR corrected). Same format as Fig. 4. (**B**) Average Z-score (±SE) across inhibitory control metrics for each subjects. (**C**) Average partial correlation of inhbotry control metrics shown in A for each subject (colors) and overall (black). The overall correlations among inhibitory control metrics was not different from zero (avg. r = 0.022 ± 0.020, p = 0.2913).

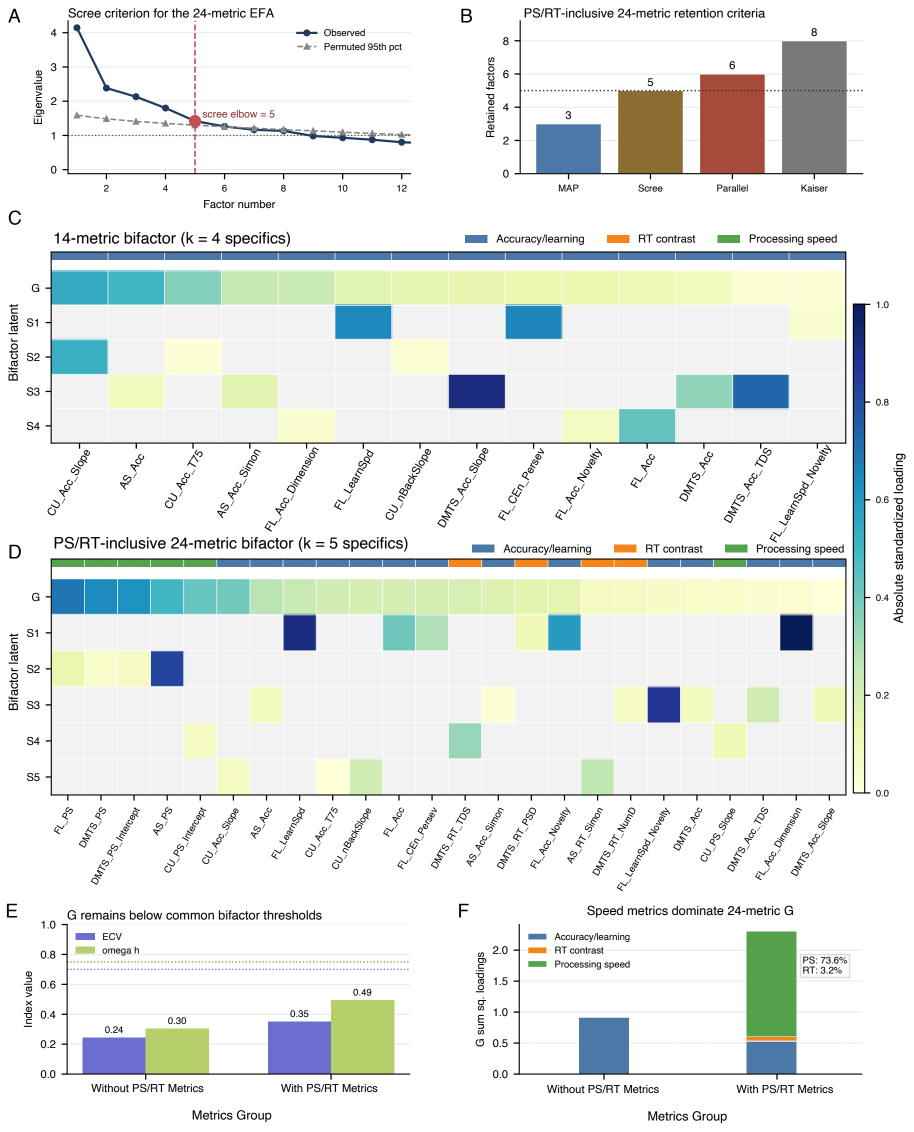


Figure S9

**Suppl. Fig. S9. Sensitivity of bifactor structure to RT contrast and processing speed metrics.** (A) Scree criterion for the speed-inclusive 24-metric EFA. The observed eigenvalue curve is shown relative to the permutation-based 95th percentile reference, with the scree elbow selecting five factors. (B) Comparison of factor-retention criteria for the 24-metric panel. MAP, scree, parallel analysis, and Kaiser criteria favored different factor counts, indicating that the speed-inclusive structure was less stable than the 14-metric accuracy-based panel. (C,D) Absolute standardized bifactor loadings for the 14-metric model and the speed-inclusive 24-metric model. The 24-metric model separates RT contrast metrics, which quantify condition-specific slowing from interference effects, from processing speed metrics, which quantify broader overall or parametric response speed. (E) General-factor interpretability indices for the two bifactor models. ECV and omega hierarchical remained below conventional thresholds for a dominant general factor. (F) Metric-class contribution to the general factor in the 24-metric model. processing speed metrics accounted for most G loading strength, whereas RT contrast metrics contributed little, suggesting that the speed-inclusive general factor primarily reflected shared response-speed variance rather than a broad common executive-function factor.

### Supplementary Tables S1-S6

**Table S1. Headline fit indices, 14-metric panel.**

| Index | Quantification |
|---|---|
| CFA(k=4) plain CFI | 0.869 |
| CFA(k=4) plain BIC | 198 |
| CFA(k=4) plain RMSEA | 0.085 |
| ESEM(k=4) CFI | 0.825 |
| ESEM(k=4) BIC | 4127 |
| BF ECV | 0.245 |
| BF omega_h(G) | 0.304 |
| HO (via S-L) ECV | 0.113 |
| HO (via S-L) omega_h(G) | 0.008 |

*4-factor CFA and ESEM fit comfortably; bifactor and higher-order G fall well below conventional reliability thresholds (ECV ≥ 0.70, $\omega$_h(G) ≥ 0.75).*

**Table S2. 14-metric panel and manuscript factor assignment.**

| Metric | Task | MS factor |
|---|---|---|
| FL_CEn_Perservation | FL | F1 (Learning) |
| FL_LearningSpeed | FL | F1 (Learning) |
| FL_LearningSpeed_Novelty_Idx | FL | F1 (Learning) |
| FL_Accuracy | FL | F3 (Exo Inh. Control) |
| FL_Accuracy_Dimension_Idx | FL | F3 (Exo Inh. Control) |
| FL_Accuracy_Novelty_Idx | FL | F3 (Exo Inh. Control) |
| DMTS_Accuracy | DMTS | F2 (Endo Inh. Control) |
| DMTS_Accuracy_Slope | DMTS | F2 (Endo Inh. Control) |
| DMTS_Accuracy_TDS_Idx | DMTS | F2 (Endo Inh. Control) |
| AS_Accuracy | AS | F2 (Endo Inh. Control) |
| AS_Accuracy_SimonCong_Idx | AS | F2 (Endo Inh. Control) |
| CU_Accuracy_Slope | CU | F4 (Memory Updating) |
| CU_Accuracy_TrialAt75Acc | CU | F4 (Memory Updating) |
| CU_nBackSlope | CU | F4 (Memory Updating) |

*MS factor name from varimax EFA dominant-loading rule (see §1 of revision plan for relabeling).*

**Table S3. Factor-count diagnostics (plain CFA + ESEM).**

| k | CFA CFI | CFA BIC | ESEM CFI | ESEM BIC |
|---|---|---|---|---|
| 1 | 0.534 | 160 | — | — |
| 2 | 0.649 | 167 | 0.678 | 4195 |
| 3 | 0.822 | 180 | 0.765 | 4142 |
| **4** | **0.869** | **198** | **0.825** | **4127** |
| 5 | 0.894 | 222 | 0.824 | 4197 |
| 6 | 0.923 | 210 | 0.724 | 4403 |

*k = 4 is the CFA elbow ($\Delta$CFI(3→4) = +0.047, plateau after). ESEM BIC bottoms at k = 4.*

**Table S4. Bifactor and higher-order reliability indices (+MTMM).**

| Factor (MS label) | BF $\omega$_s | BF H | BF FD | HO $\omega$_s | HO $\gamma$→G |
|---|---|---|---|---|---|
| G (general) | 0.304 | 0.535 | 0.738 | 0.008 | — |
| F1 Flex Learn | 0.446 | 0.593 | 0.770 | 0.000 | 0.504 |
| F2 Endo Inh | 0.038 | 0.872 | 0.938 | 0.127 | -0.262 |
| F3 Exo Inh | 0.073 | 0.187 | 0.433 | 0.241 | 0.258 |
| F4 WM Update | 0.070 | 0.275 | 0.561 | 0.197 | 0.351 |

*BF and HO G factor and specifics fail conventional reliability thresholds ($\omega$_s ≥ 0.50 substantive; $\omega$_h(G) ≥ 0.75 reliable).*

**Table S5. Replication and cluster discrimination (CF model).**

| Factor (MS) | Split-half median $\phi$ | LOSO median $\phi$ | LOSO min $\phi$ | Cluster $\eta^2$ | p |
|---|---|---|---|---|---|
| F1 Flex Learn | 0.962 | 0.947 | 0.739 | 0.271 | 0.623 |
| F2 Endo Inh | 0.926 | 0.961 | 0.731 | 0.376 | 0.493 |
| F3 Exo Inh | 0.887 | 0.965 | 0.911 | 0.328 | 0.551 |
| F4 WM Update | 0.952 | 0.982 | 0.797 | 0.890 | 0.037 |

*All 4 factors replicate at median $\phi$ ≥ 0.88. Factor 4 (WM Updating) is the only factor with significant cluster discrimination.*

**Table S6. Bootstrap 95% CIs on G-factor indices (B = 60).**

| Index | Point | CI 2.5% | CI 97.5% |
|---|---|---|---|
| BF ECV | 0.245 | 0.159 | 0.536 |
| BF $\omega$_h(G) | 0.304 | 0.002 | 0.435 |
| BF $\omega$_total | 0.437 | 0.180 | 0.608 |
| HO ECV (S-L) | 0.113 | 0.142 | 0.563 |
| HO $\omega$_h(G) (S-L) | 0.008 | 0.005 | 0.433 |
| $\gamma$ → F1 Flex Learn | 0.504 | 0.087 | 1.000 |
| $\gamma$ → F2 Endo Inh | -0.262 | -1.000 | 1.000 |
| $\gamma$ → F3 Exo Inh | 0.258 | -0.330 | 1.000 |
| $\gamma$ → F4 WM Update | 0.351 | -0.630 | 1.000 |

*CIs span both unity and diversity regimes for BF; HO $\gamma$ paths show sign flips. Both confirm G is not stable.*

**Suppl. Table ST1. Summary of model-validation, factor-assignment, and general-factor diagnostics for the 14-metric panel.**

(S1) Headline model indices for the 14-metric panel, including plain four-factor CFA fit, four-factor ESEM fit, bifactor general-factor indices, and higher-order general-factor indices. (S2) Retained metrics, task origin, and manuscript factor assignment for the 14-metric panel. (S3) Factor-count diagnostics from plain CFA and ESEM across alternative factor numbers, showing the four-factor solution as the CFA elbow and the ESEM BIC minimum. (S4) Bifactor and higher-order reliability indices for the general and specific factors, showing weak reliability for a unitary general factor. (S5) Split-half and leave-one-subject-out factor congruence, together with cluster-discrimination statistics for each manuscript factor. (S6) Bootstrap confidence intervals for bifactor and higher-order general-factor indices. Together, these diagnostics support a four-factor correlated structure while arguing against a stable, reliable unitary general factor in the present dataset.

**Supplementary Movie S1: Example performance of trials of the Antisaccade task of three subjects.** The subject sits in his home cage, one arm touching the monitor center where the start button appears, and the other hand holding on to the sipper tube that will release fluid. The display background of the AS task had a structured blueish sky theme. A token bar with eight placeholder tokens is presented on top of the screen. Three token placeholders are green at the start of the trials reflecting the starting assets. Subjects touch a blue circular start-button to initiate a trial. Then a cue (grey asterisk) and a uniform colored grey Quaddle with straight arms appear left or right of the screen, followed by a display of four Quaddle objects in the response display. The target can be one of four different body shapes (hourglass, oval cylindrical, cube) and the response displays shows all four shapes. The subject choses the target shape in the response display. Three reward tokens appear above the chosen object and move towards the token bar where they are added as green tokens in empty (gold-colored) placeholders token positions When the subject has 6 of 8 tokens, the subjects receives two final tokens for a correct choice, which complete the eight token place holders, causing a red/white flashing of the token bar and the delivery of three reward pulses. In the video the three reward pulses are heard in the background, indicating that the LMI reward pump pushed water through the sipper tube that is in front of the subject.

**Supplementary Movie S2: Example performance of trials of the Continuous Updating task.** The subject initiates a block which starts in trial one by displaying two objects. Choosing (touching) one object triggers a yellow halo around the chosen object and increments a green slider position and triggers immediate fluid reward. The second trial shows both previous and one new object at random locations of a virtual 9x6 grid spanning the monitor. Successive correctly performed trials adds new objects until the subject choses an object that was previously already chosen. This error is followed by a grey halo around the chosen object, the green slider position retracts to the left starting position of the slider, and the new block starts by displaying the start button. Once the start button is touched, two new objects appear in the first trial of the new block and the subject continues performing the block until he makes an error. Each session allowed performing 50 blocks.

**Supplementary Movie S3: Example performance of trials of the Flexible Learning task.** The subject initiates a trial by touching a blue start button, three objects appear at quasi-random locations on the circumference of a virtual circles equidistant from the monitor center. The subject than chooses one of the three objects, receives visual feedback (a yellow halo for correct and a grey halo for choosing a rewarded / unrewarded object), followed by tokens appearing on top of the choice-location. A token bar with 8 place holder tokens is presented on top of the screen. The tokenbar contains 3 green starting token. When the subject makes an error (the first tiral in the video) the subject sees a grey halo and grey visual tokens that are animated towards to the token bar whether they 'substract' three green tokens, which disappear, indicating the loss of 3 already attained tokens. The task involved losing 1 or 3 tokens for incorrect choices and gaining2 or 3 tokens for correct choices in different blocks of trials. Subjects learned from trial and error which object feature is rewarded when chosen. The rewarded feature could be from any of six different features from two feature dimensions, e.g. the first trials of the video shows obejcts varying in color and body shape and only the object with the square body shape will be rewarded. Te background image is randomly picked from a sand thema (see text for details).

**Supplementary Movie S4: Example performance of trials of the Delayed Match-to-Sample task.** The subject initiates a trial by touching a blue start button. The sample stimulus appears at a random location followed by a variable delay of 0.3-5.1 s. Then three or four objects appear at random locations and the subject has to choose (touch) the object that matches the sample. The objects are Quaddles that vary in three dimensions. In the video the trial shows objects that vary in arm style, color, and shape, while the second trial has objects carrying in color, shape, and surface patterns. The context background contains a random image with rocks from a rock thema that we specifically associate with the delayed match-to-sample task (see Methods). The top of the screen shows the token bar. When subjects correctly chose the sample object in the response display he received a yellow halo around the chosen object as visual feedback, and two green tokens appear above the chosen object and are animated to move up and into the token bar. The token bar starts empty and correct choices add two tokens initially and the remaining 3$^{rd}$ token after the second correct choice. When the token-bar fills, it flashed red/white and two fluid pump openings provide water through the sipper tube.
